# The 5’-NAD cap of RNAIII modulates toxin production in *Staphylococcus aureus* isolates

**DOI:** 10.1101/778233

**Authors:** Hector Gabriel Morales-Filloy, Yaqing Zhang, Gabriele Nübel, Shilpa Elizabeth George, Natalya Korn, Christiane Wolz, Andres Jäschke

## Abstract

1

Nicotinamide adenosine dinucleotide (NAD) has been found to be covalently attached to the 5’-ends of specific RNAs in many different organisms, but the physiological consequences of this modification are largely unknown. Here we report the occurrence of several NAD-RNAs in the opportunistic human pathogen *Staphylococcus aureus*. Most prominently, RNAIII, a central quorum-sensing regulator of this bacterium’s physiology, was found to be 5’-NAD-capped to a significant extent. NAD incorporation efficiency into RNAIII was found to depend *in vivo* on the −1 position of the P3 promoter. Reduction of RNAIII’s NAD content led to a decreased expression of alpha- and delta-toxins, resulting in reduced cytotoxicity of the modified strains. These effects to not seem to be due to changes in RNAIII’s secondary structure upon NAD attachment, as indicated by largely unaltered patterns in *in vitro* chemical probing experiments. Our study represents a large step towards establishing a biological function of the 5’-NAD cap, which for RNAIII in *S. aureus* is to modulate the expression of virulence factors.

**Importance:** Numerous organisms, including bacteria, are endowed with a 5’-NAD cap in specific RNAs. While the presence of the 5’-NAD cap modulates the stability of the modified RNA species, a significant biological function and phenotype have not been assigned so far. Here, we show the presence of a 5’-NAD cap in RNAIII from *S. aureus,* a dual-function regulatory RNA involved in quorum-sensing processes and regulation of virulence factor expression. We also demonstrate that altering the natural NAD modification ratio of RNAIII leads to a decrease in exotoxin production, thereby modulating bacterium’s virulence. Our work unveils a new layer of regulation of RNAIII and the *agr* system that might be linked to the redox state of the NAD molecule in the cell.

## 3 Introduction

The discovery of the nicotinamide adenosine dinucleotide (NAD) bacterial 5’-cap in regulatory RNAs in *Escherichia* coli (Cahová, Winz, Höfer, Nübel, & Jäschke, 2015) challenged a long-standing dogma. To date, the 5’-NAD cap has been reported in Gram-negative and Gram-positive (*Bacillus subtilis* (Frindert et al., 2018)) bacteria as well as in eukaryotes such as *Saccharomyces cerevisiae* (Kellner et al., 2014; Walters et al., 2017), the mammalian cell line HEK293T (Jiao et al., 2017) and plants from the genus *Arabidopsis* (Wang et al., 2019; Zhang et al., 2019). This novel nucleotide modification in RNA appears to be present in most organisms if not ubiquitous. However, no NAD-modified RNA has been reported in pathogens yet. The opportunistic pathogen *Staphylococcus aureus* is a gram-positive bacterium responsible for the majority of nosocomial infections (David & Daum, 2010). After the invasion of host tissue, *S. aureus* can cause bacteremia, sepsis, endocarditis, and toxic shock syndrome (Lowy, 1998). The pathogenicity of *S. aureus* is strongly related to the expression of numerous virulence factors, many of which are under the control of the *agr* quorum-sensing system (Bronesky et al., 2016; Quave et al., 2016; Traber et al., 2008). The *agr* locus harbours two promoters (P2 and P3) located back to back in the chromosome (Janzon, Lofdahl, & Arvidson, 1989; Peng, Novick, Kreiswirth, Kornblum, & Schlievert, 1988) (Figure 1A). P3 encodes a regulatory RNA (RNAIII) that additionally contains a small open reading frame (ORF) for delta-toxin (Hld) (Janzon & Arvidson, 1990; Novick et al., 1993), whereas P2 regulates the transcription of a polycistronic RNA (RNAII) that encodes the Agr proteins Agr A/B/C/D (Figure 1A) (Novick et al., 1995). With this quorum-sensing mechanism, *S. aureus* senses the cell density based on the extracellular concentration of the self-produced autoinducer peptide (AIP) (Recsei et al., 1986). The AIP binds to the AgrC histidine kinase membrane receptor and activates AgrA regulator by phosphorylation (Figure 1A). AgrA∼P activates transcription of the P2 and P3 promoters, producing RNAII and RNAIII, respectively (Figure 1A) (Bronesky et al., 2016; Koenig, Ray, Maleki, Smeltzer, & Hurlburt, 2004). AgrA∼P furthermore activates transcription of phenol-soluble modulins (*psmα* and *psmβ*) (Queck et al., 2008), which are small peptides with surfactant activity that contribute to cell membrane disruption and therefore to *S. aureus* cytotoxicity (M. Li et al., 2009; Peschel & Otto, 2013). RNAIII plays a crucial bifunctional role in the switch from the adhesion phase towards invasion (Bronesky et al., 2016; Koenig et al., 2004), which ultimately determines the infective behaviour of the bacterium. On the one hand, it inhibits translation of *rot* mRNA, which encodes for a repressor (Rot) that blocks the transcription of several toxins (Boisset et al., 2007; Geisinger, Adhikari, Jin, Ross, & Novick, 2006; Said-Salim et al., 2003). Also, RNAIII represses the expression of surface proteins, e.g., protein A or coagulase through an antisense mechanism (Boisset et al., 2007; Bronesky et al., 2016; Huntzinger et al., 2005). On the other hand, it activates the production of several hemolytic toxins, such as alpha-toxin (Hla) (Morfeldt, Taylor, von Gabain, & Arvidson, 1995) and the self-encoded Hld.

**Figure 1:**
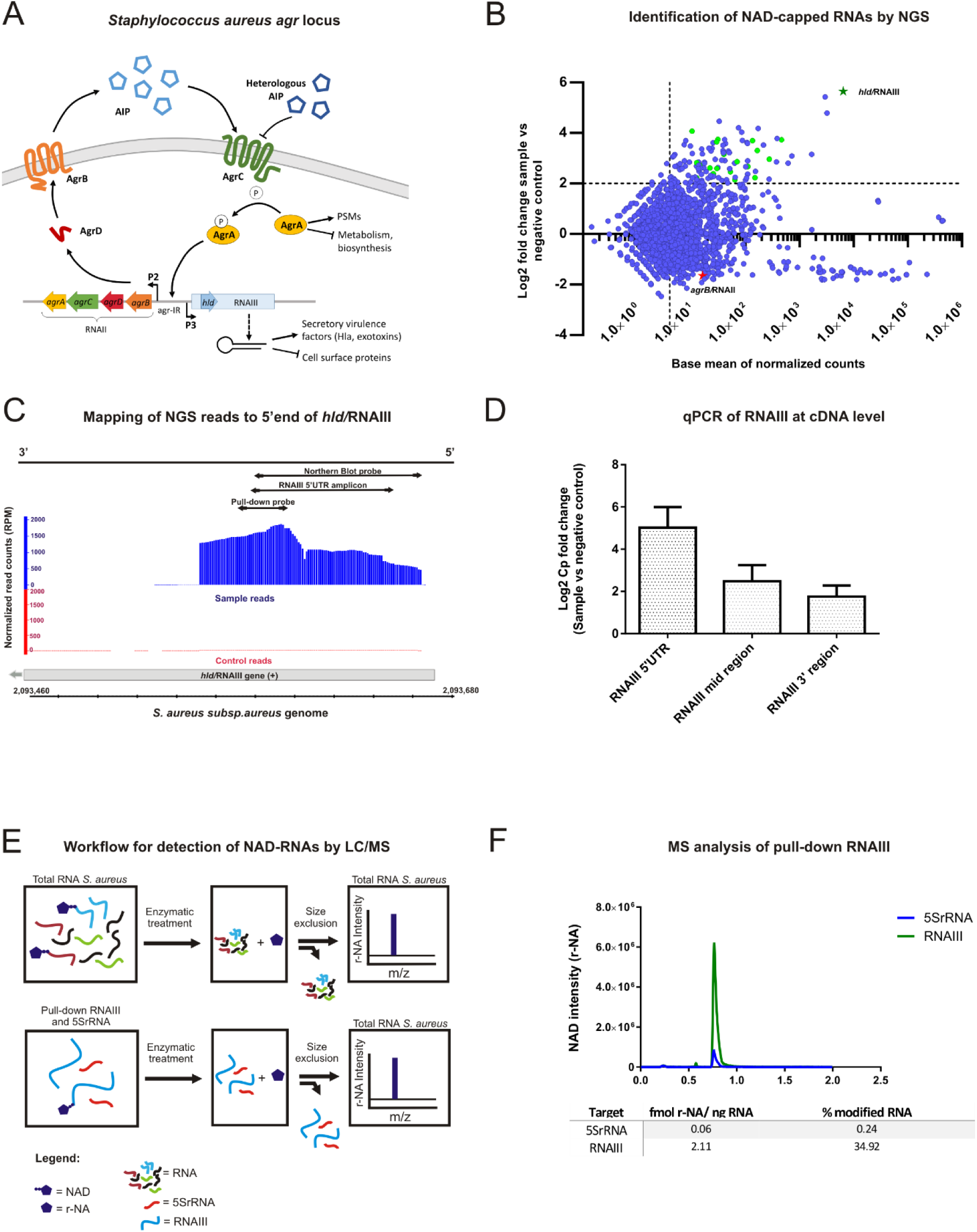
**A**: Schematic representation of the cell-population-sensor Agr locus in *S. aureus*. AgrD is a precursor peptide that is converted into an autoinducing peptide (AIP) upon proteolysis by AgrB (transmembrane protein) and secreted to the exterior. The AIP then binds to the AgrC receptor, triggering autophosphorylation of its intracellular histidine kinase domain. AgrA is activated by the transfer of the phosphate group of AgrC and enhances transcription of P2 and P3 promoters, leading to RNAII and RNAIII production. RNAII encodes all the Agr proteins (AgrB/D/C/A) whereas RNAIII is a regulatory RNA also containing an ORF encoding for delta-toxin (Hld) (See text for more details). **B**: Scatter plot of NGS data after NAD captureSeq. Y-axis represents the enrichment of RNAs (Log2 Fold change) in the fully treated samples against minus ADPRC (control). The X-axis shows the average of normalized counts. The dots confined in the upper right region of the plot represent the RNAs significantly enriched (NAD-capped RNAs). The light green dots depict the hits clustering at the 5’-UTR and with +1A. A light green star represents the *hld*/RNAIII gene, which is the most enriched by far, whereas RNAII (depicted by a red star) is located in the non-enriched area of the graph. **C:** Distribution of normalized reads on the *hld*/RNAIII gene (grey bar) visualized with the Integrated Genome Browser (Nicol et al., 2009). The sample reads (shown in blue) are the normalized reads of the ADPRC-treated sample group whereas the red-labeled control reads are the ADPRC-negative. The orientation of the gene is indicated in the black bar above and represents the coding strand (+). RPM: Reads per million mapped reads. **D:** Bar chart representing the enrichment at the cDNA level of three targeted regions of the hld/RNAIII gene (5’-UTR, middle region, and 3’-region). The Y-axis represents the Cp fold change (sample vs. negative control group) obtained by qPCR. **E:** Schematic view of workflow followed for detection of NAD-RNAs in total RNA samples and pulled-down RNAs. **F:** Extracted ion chromatogram representing the pull-down LC-MS outcome. The Y-axis represents the NAD (r-NA) intensity of RNAIII (green, from 40 ng RNA) and 5SrRNA (blue, negative control, from 960 ng RNA). The X-axis indicates the retention time in minutes. The table beneath shows the concentration of r-NA (NAD) per ng RNA and also the NAD modification percentage for each RNA.

The addition of the canonical N7-methyl guanosine (m^7^G) cap to the 5’-ends of eukaryotic mRNAs proceeds co-transcriptionally once the transcript has reached a length of ∼25 nucleotides. The NAD cap is, however, incorporated in a different way in prokaryotes and probably in all organisms: the ubiquitous redox coenzyme NAD is incorporated as the very first nucleotide (*ab initio*) into RNA during transcription initiation by the RNA polymerase, acting as a non-canonical initiating nucleotide (NCIN) that competes with ATP at A-initiating promoters (Bird et al., 2016; Frindert et al., 2018). Thus, transcription appears to constitute the primary source of 5’-NAD-RNAs in the cell. NAD incorporation into RNA has furthermore been demonstrated to strongly depend on the promoter sequence around the transcription start site (TSS) (Frindert et al., 2018; Vvedenskaya et al., 2018). In *E. coli* and *B. subtilis,* the −1 position turned out to be the most important one for NAD incorporation, likely due to pseudo-base-pairing between the nicotinamide moiety of NAD and the nucleotides at the −1 position of the DNA non-coding strand (thymidine (T) and cytidine (C)) (Vvedenskaya et al., 2018).

The functions of the NAD cap are not yet fully understood, but in prokaryotes, it seems to confer stability to RNA. In *E. coli,* the NAD cap protects RNA against 5′-processing by the RNA-pyrophosphohydrolase (RppH) and thereby against RNaseE degradation (Cahová et al., 2015), whereas in *B. subtilis*, it stabilises RNA against exoribonucleolytic attack by RNase J1 (Frindert et al., 2018). Furthermore, *E. coli* possesses an NAD-decapping machinery based on the Nudix phosphohydrolase NudC (Frick & Bessman, 1995), which efficiently hydrolyses the NAD cap and leaves a 5’-monophosphate RNA (p-RNA) (Cahová et al., 2015). Despite Nudix hydrolase motifs being present in several enzymes of *S. aureus,* no NudC ortholog has been found to date. In mammalian cells, the NAD-decapping machinery works in a very different way. The DXO enzymes first remove the entire NAD cap from an mRNA (releasing NAD and p-RNA) and then proceed with exonucleolytic mRNA decay (Jiao et al., 2017).

In this study, we reveal by NAD captureSeq the presence of NAD-capped mRNAs and regulatory RNAs in the opportunistic pathogen *S. aureus.* The highly expressed, multifunctional, protein-coding, and regulatory RNA, RNAIII, is found to bear the NAD cap. Additionally, we study the importance of the −1 and +1 positions of the P3 promoter for the incorporation of NAD into RNAIII *in vivo*. Finally, we also investigate the consequences of changing RNAIII’s NAD modification ratio on *S. aureus* virulence. The obtained results are interpreted in the context of the current knowledge of *S. aureus’s* pathobiology.

## 4 Results

### 4.1 *S. aureus* possesses NAD-capped RNAs

To detect and quantify NAD-capped RNAs in *S. aureus,* Liquid Chromatography coupled with Mass Spectrometry (LC-MS) analysis of total RNA samples was carried out. Two RNA samples from *S. aureus* ATCC 25923 were washed extensively to remove contaminating free NAD before enzymatic treatment with *E. coli* NudC and alkaline phosphatase. Upon this treatment, NAD-RNAs will liberate N-ribosylnicotinamide (r-NA) that can be sensitively detected by MS analysis. The results revealed an average amount of 25.25 ± 1.64 fmol of NAD-RNA per microgram of RNA, which corresponds to an estimated amount of 897 ± 58 NAD-RNA molecues per cell (Table S1) (Chen, Kowtoniuk, Agarwal, Shen, & Liu, 2009). These results indicated the presence of NAD-capped RNAs in *S. aureus*.

In order to identify NAD-capped RNAs, NAD captureSeq was performed (Winz et al., 2017) Total RNA from *S. aureus* ATCC 25923 (isolated at the late exponential phase) was treated per duplicate either with ADP-ribosyl cyclase (ADPRC), allowing selective biotinylation of NAD-RNA, or mock-treated (minus ADPRC control). The captured RNAs were reverse transcribed and PCR-amplified to generate a DNA library for Illumina sequencing (Figure S1A). The distribution of the obtained normalised mapped reads revealed 96.87% mRNAs, 2.46% tRNAs, and 0.68% rRNAs with enrichment values up to 50-fold. Enriched hits (*P* < 0.05, log2-fold change > 2, base mean > 10) were visualised with the Integrated Genome Browser (Nicol, Helt, Blanchard, Raja, & Loraine, 2009) to identify reads that clustered at the 5’-ends of transcripts initiating with adenosine (+1A) (Figure 1B). 67.7% of the initial enriched hits (21 out of 31) had these properties (Table S2) and are indicated as green dots in the upper right sector of the scatter plot (Figure 1B). The TSS analysis of the enriched hits showed T (61.9%) or A (28.6%) at position −1 of the promoter predominantly. G or T usually occupy the +2 position with the same occurrence (33.3%) whereas +3 position is dominated by A (57.1%). The predominant nucleobase at position -3 and -2 of the promoter was A (71.4% and 47.6% respectively, Figure S1B). Most enriched RNAs turned out to be mRNA 5’-fragments (Table S2). However, the most abundant and the most enriched hit was the bifunctional RNAIII (*hld*/RNAIII gene, Figure 1B, green star). Notably, the other transcript from the agr locus, RNAII, which has a similar promoter and is reported to also initiate with A (Novick et al., 1995; Reynolds & Wigneshweraraj, 2011), was not enriched at all (Figure 1B, red star).

### 4.2 The multifaceted regulatory RNAIII contains a 5’-NAD cap

The obtained reads for *hld/*RNAIII gene mapped clearly on its 5’-UTR (Figure 1C). As NAD captureSeq does not use a fragmentation step and size-selects (at the very end, after PCR amplification) amplicons that correspond to RNAs < ∼170 nt, information about full-length mRNA species cannot be obtained from these data. To quantitatively investigate the length distribution of NAD-RNAIII, qPCR was performed on the enriched cDNA obtained directly from reverse transcription, before any amplification or size fractionation using gene-specific primers (Table S3) targeting different regions of RNAIII, thereby comparing the cDNA from the NAD captureSeq sample with the mock-treated negative control. The qPCR data revealed the most substantial enrichment for the 5’-end region, but a significant presence and enrichment of full-length RNAIII (Figure 1D). Northern blot analysis (which is independent of the nature of the 5’-end) indicated that overall, RNAIII in *S. aureus* is predominantly full-length (Figure S1C).

To confirm that the NAD is covalently linked to RNAIII’s 5’-end, RNAIII (enriched) and 5SrRNA (non-enriched, negative control) were specifically isolated from *S. aureus* total RNA by pull-down using biotinylated complementary oligonucleotides. Afterwards, RNAs were quantified, treated with NudC and alkaline phosphatase, and the small-molecule fraction analysed by LC-MS (Figure 1E). This analysis revealed an intensive peak corresponding to nicotinamide (m/z = 122.81, originating from r-NA, m/z = 254.94) in the case of RNAIII, whereas for 5SRNA the same peak could only be detected at 24-fold higher input RNA concentration (Figure 1F). The NAD modification ratio, i.e., the percentage of molecules of that RNA species that carry the 5’-NAD-modification, of RNAIII and 5SrRNA was calculated as 36.20% and 0.25%, respectively (Figure 1F).

### 4.3 Guanosine at position −1 of the P3 promoter increases NAD incorporation into RNAIII in vivo

To investigate the biological consequences of 5’-NAD modification in vivo, *S. aureus* strains that differ strongly in their content of 5’-NAD-RNAIII, with as little as possible differences in the transcriptome, had to be generated. Toward this end, we complemented a strain devoid of the *hld/*RNAIII gene (*S. aureus* HG001 *ΔRNAIII*) with plasmids based on the shuttle vector pCG-246 (Helle et al., 2011). These plasmids carried the hld/RNAIII gene behind different promoters: pCG-P3 carried the native P3 promotor, while pCG-P3(−1G) had a mutated version of this promoter (−1T to −1G). To unravel whether the lack of NAD captureSeq enrichment of RNAII was due to properties of its promoter, construct pCG-P2 was generated in which the P2 promoter controlled the RNAIII native sequence (Figure 2A). Unlike the previously used *S. aureus* ATCC 25923, in the generated strains the *hld/*RNAIII gene is located on a plasmid.

**Figure 2:**
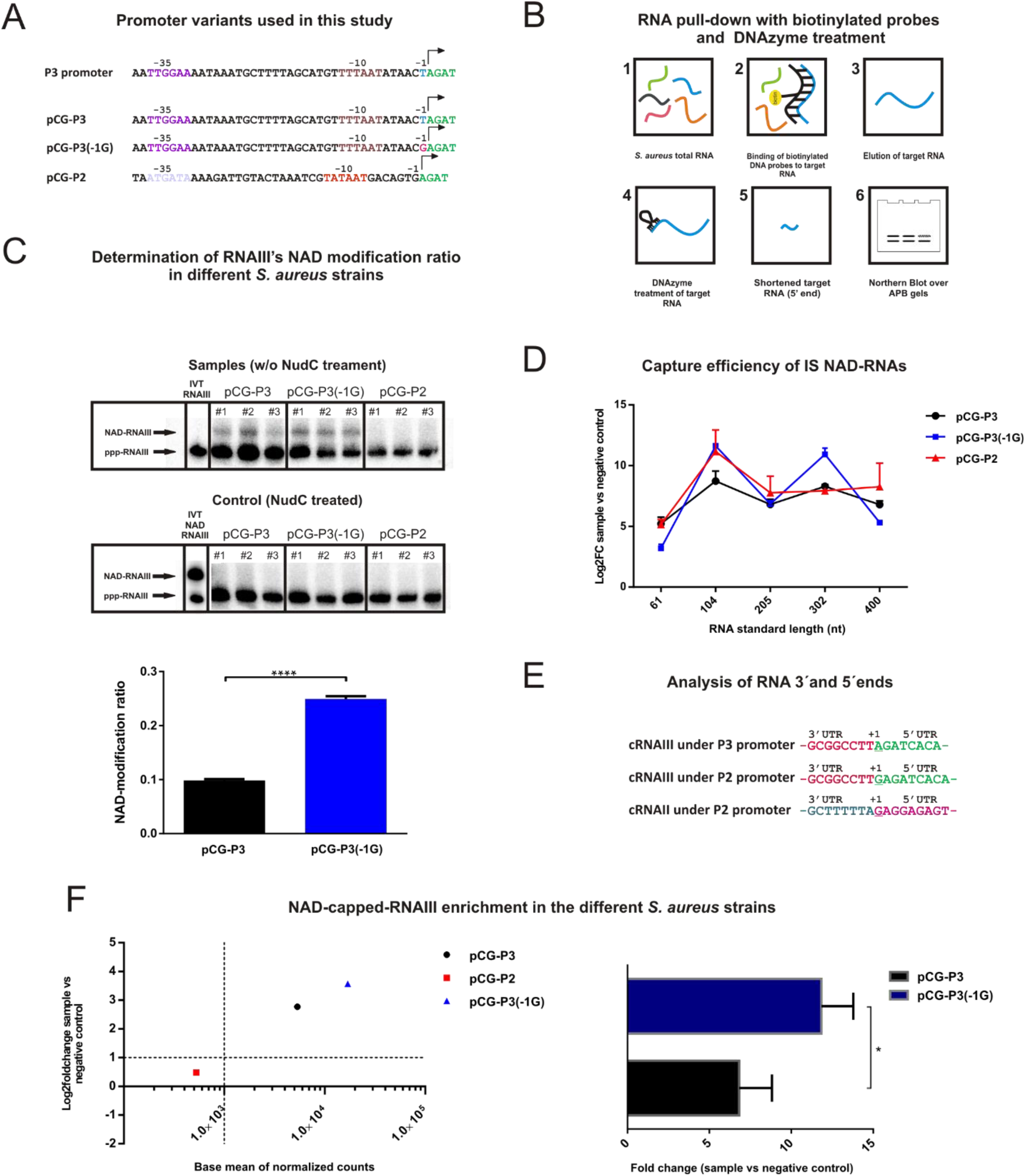
**A:** Promoter sequences of the different pCG-246-derivated constructs used in the study (pCG-P3, pCG-P3(−1G), and pCG-P2). Highlighted are the -35 (-35, purple for P3 and light gray for P2), −10 (−10, brown for P3 and orange for P2) and −1 (−1, light blue for native P3, pink for mutated P3 and black for native P2) regions. The first four nucleotides of RNAIII are highlighted in green. **B:** The figure shows the workflow (steps 1 to 6) followed for the pull-down of specific RNA targets and the subsequent RNA cleavage by a DNAzyme (see text for more information). **C:** Analysis of pulled-down RNAIII by Northern blot on APBgels after DNAzyme treatment of the different *S. aureus* strains (pCG-P3, pCG-P3(−1G), and pCG-P2, each of them by triplicate). The arrows indicate the bands belonging to NAD-RNAIII and ppp-RNAIII 5’-termini. As a marker, 10 ng of *in vitro*-transcribed (IVT) ppp-RNAIII treated with DNAzyme were loaded in the first lane from the left. Control samples underwent a NudC treatment before analysis on APBgels. The marker used was mixed NAD/ppp-RNAIII instead of ppp-RNAIII. The bar chart beneath depicts the quantification of NAD-modification ratio of RNAIII of *S. aureus* pCG-P3 (black) and pCG-P3(−1G) (blue), based on the Northern blot without NudC treatment. Statistical significance determined by *t-*test, n = 5. **D:** The chart represents the enrichment (capture efficiency) of each of the IS NAD-RNAs in the three NAD captureSeq experiments conducted: *S aureus* pCG-P3 (black dot and line), pCG-P3(−1G) (blue square and line) and pCG-P2 (red triangle and line). **E:** Scatter plot of NGS data after NAD captureSeq. Y-axis represents the enrichment of *hld/*RNAIII gene (log2fold change) in the fully treated samples against minus ADPRC (control). The X-axis shows the average of normalised counts. Each strain is represented with markers: a black dot for pCG-P3, blue triangle for pCG-P3(−1G), and red square for pCG-P2. The bar chart on the right side shows the RNAIII enrichment in terms of fold change (sample vs negative control) of pCG-P3 (black bar) and pCG-P3(−1G) (blue bar) *S. aureus* strains. Statistical significance determined by *t-* test, n = 6. **F:** The figure shows the ligated 3’-UTR and 5’-UTR of circularised RNAs (cRNA). Sequences were obtained by Sanger sequencing after cRT-PCR (see Methods). *P* < 0.05; *, *P* < 0.0001; ****. Horizontal bars represent mean. Error bars depict standard deviation.

To detect variations in RNAIII’s modification ratio, we analysed RNAIII pulled down from total RNA from the RNAIII-complemented *S. aureus* strains by acryloaminophenyl boronic acid (APB) gel electrophoresis that separates 5’-NAD-RNA from 5’-p and 5’-ppp-RNA, combined with Northern blot (Alwine, Kemp, & Stark, 1977; Igloi & Kossel, 1985; Nübel, Sorgenfrei, & Jäschke, 2017). As the full-length RNAIII was too long for efficient separation and precise quantification (Figure S1D), a pre-treatment with a designed DNAzyme was introduced which cleaved RNAIII to yield 125 nt 5’-terminal fragments (Figure 2B). The strains containing pCG-P2 showed no NAD incorporation into the RNAIII transcript at all, whereas RNAIII in pCG-P3 and pCG-P3(−1G)-transformed strains showed an average NAD-modification ratio of 9.82± 0.15% and 24.91 ± 0.37%, respectively (Figure 2C). Indeed, *S. aureus* pCG-P(−1G) was found to have a significantly higher amount of NAD-RNAIII than *S. aureus* pCG-P3 (*t*-test; *P* < 0.0001), demonstrating the importance of the −1 mutation on NAD incorporation efficiency, which confirms findings in other microorganisms (Frindert et al., 2018; Vvedenskaya et al., 2018). In all RNAIII samples, the slower-migrating band disappeared upon treatment with *E. coli* NudC, further supporting the nature of the 5’-modification as NAD (Figure 2C).

To test whether this different NAD modification percentage in the three RNAIII constructs (pCG-P3, pCG-P2, pCG-P3(−1G)) is also reflected in NAD captureSeq enrichment, we had to modify the analysis to allow quantitative comparisons between different RNA species by adding pure NAD-RNA spike-in controls of five different lengths as internal standards (IS; 61, 104, 205, 302 and 400 nt, Table S4) to allow for normalization. The NGS results revealed that the NAD capture efficiency of the IS between the samples was very similar (Figure 2D, one-way ANOVA; *P* = 0.8617). When comparing the *hld*/RNAIII enrichment in the different *S. aureus* strains, pCG-P3(−1G) (log2FoldChange = 3.57) showed higher enrichment than *S. aureus* pCG-P3 (log2FoldChange = 2.77). In the case of *S. aureus* pCG-P2, no enrichment was found for *hld*/RNAIII (log2FoldChange = 0.48). Thus, the NGS data support the results obtained by APBgel - Northern blot analysis and confirm that a −1G in the P3 promoter significantly enhances NAD incorporation into RNAIII (*t-* test; *P* = 0.036, Figure 2E). Analysis of the NAD captureSeq data from the three mutant strains revealed a similar pattern of NADylation as in the ATCC 25923 wild type strain (Table S5, Table S6 and Table S7). For pCG-P3 we obtained 11 enriched hits (4 of them common with the wild type), in pCG-P3(−1G) 12 (4 commons), and pCG-P2 6 (3 commons).

### 4.4 The TSS of the P2 promoter is guanosine

We were puzzled by the observation that RNAIII from strain pCG-P2 seemed to contain no NAD at all. Sanger sequencing data after cRT-PCR (Slomovic & Schuster, 2013), however, provided a straightforward explanation: the transcription start site of the P2 promoter was a G instead of the previously reported A (Novick et al., 1995; Reynolds & Wigneshweraraj, 2011), and NAD does not compete with GTP in transcription initiation. The analysis of 96 sequencing reactions (96 single colonies from each strain) from *S. aureus* pCG-P3 and pCG-P2 yielded well-defined 3’ and 5’ ends of RNAIII, where the RNAIII’s first nucleotide was +1A when transcribed from the P3 promoter, and +1G in the case of the P2 promoter (Figure 2F). A +1G was also found in RNAII (a natural product of P2 promoter) from *S. aureus* HG001 wild type strain after the analysis of 10 Sanger sequencing reactions (10 single colonies, Figure 2F).

### 4.5 The 5’-NAD cap of RNAIII affects Hla and Hld expression levels and thereby *S. aureus’s* cytotoxicity

To study the accessibility of its NAD-5’-end, ^32^P-body-labelled pure NAD-RNAIII was subjected to *in vitro* cleavage with *E. coli* NudC. After 20 min of incubation at 37 °C, NudC had decapped ∼65% of the RNAs, and this value did not change upon further incubation (up to 1 hour, Figure 3A). An additional unfolding-folding cycle, followed by the addition of fresh NudC, increased decapping to ∼83% (Figure 3A), suggesting that a fraction of NAD-RNAIII exists in a structure that is not susceptible to NudC cleavage, e.g., with the 5’-end involved in a double-strand (Höfer, Abele, Schlotthauer, & Jäschke, 2016).

**Figure 3:**
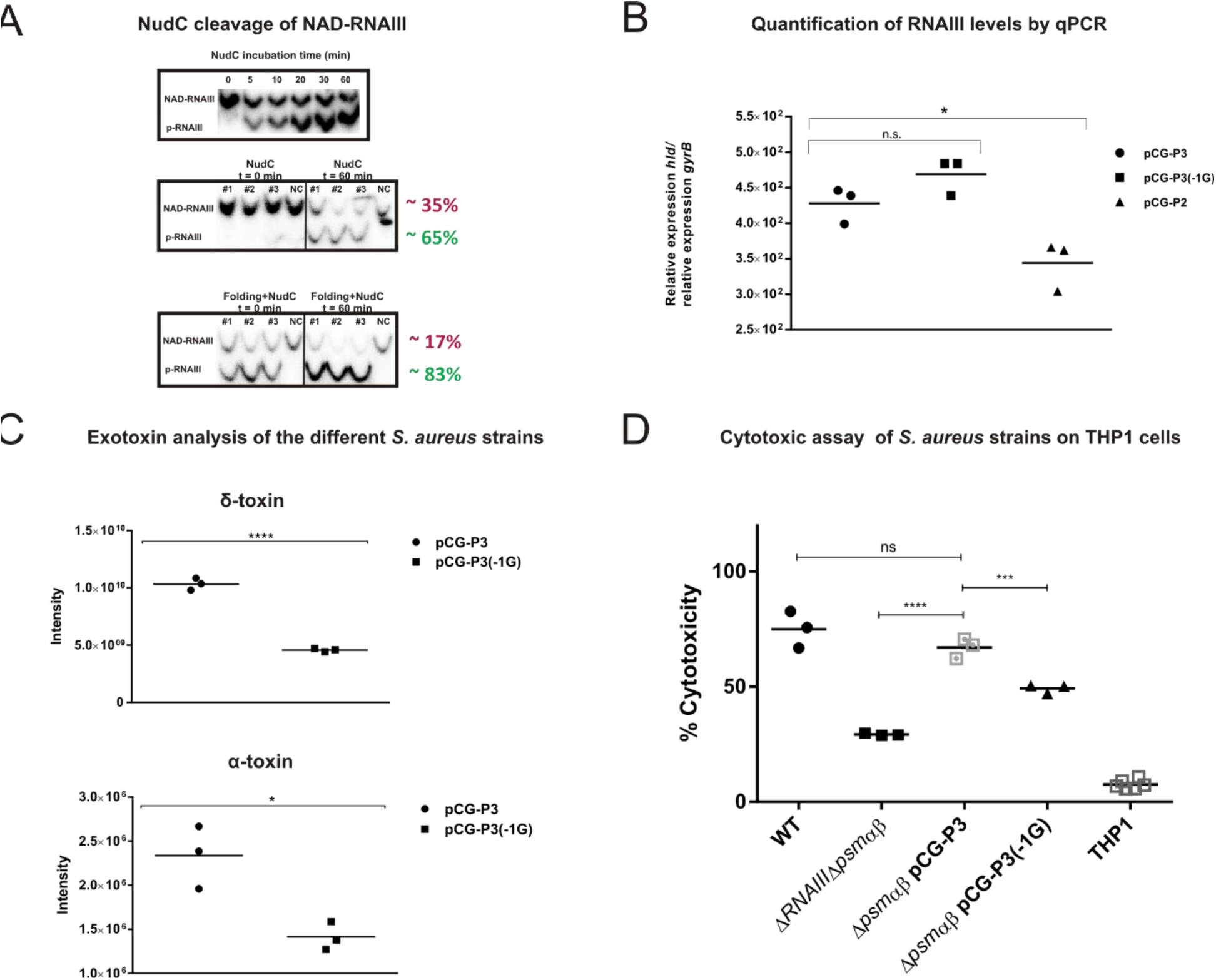
A: Cleavage of NAD-RNAIII by NudC. The reactions were analysed on APBgels. The bands represent radiolabeled (body labelled) NAD-RNAIII and p-RNAIII. Incubation controls withouth enzyme (NC) were included. #1, #2, and #3 depict the three replicates. Highlighted in red and green are the percentages of NAD-RNAIII and pRNA after incubation. RNA folding plus the addition of fresh NudC afterwards was done to equilibrate the RNAIII secondary structure pool again. **B:** RNAIII levels in the three different strains measured by qPCR. Y-axis represents the relative expression of *hld* divided by the relative expression of the housekeeping gene *gyrB.* Statistical significance determined by *t-*test, n = 6. **C:** The two scatter plots show the total amount of Hld and Hla in the supernatant of *S. aureus* pCG-P3 and pCG-P3(−1G) strains. The Hld intensity was calculated by integration of the two highest MS peaks: Hld m/z = 993.5 and its formylated version m/z = 1002.9. Alpha-toxin was detected by Western blotting and quantified by densitometry. Statistical significance determined by *t-*test, n = 6. **D:** Bacterial supernatants were analysed for cytotoxic potential against THP1 macrophages. Percent cytotoxicity shown was normalised to the Triton control. Statistical significance determined by one-way ANOVA with Tukey’s post-test, n = 21. \**P* ≤ 0.05, \*\**P* ≤ 0.01, \*\*\**P* ≤ 0.001, \*\*\*\**P* ≤ 0.0001, ns: non-significant. WT: wild type *S. aureus* HG001. Horizontal bars represent mean.

To test the effect of the promoter mutations on the biosynthesis rate of RNAIII, a qPCR assay was performed with cDNA from the different *S. aureus* strains (total RNA harvested in late exponential phase and reverse transcribed). The assay revealed no significant difference in RNAIII content between pCG-P3 and pCG-P3(−1G) strains (*t-*test; *P* = 0.1221, Figure 3B). On the other hand, pCG-P2 showed a significantly lower amount of RNAIII than pCG-P3 (*t-*test; *P* = 0.0276, Figure 3B).

Strains pCG-P3 and P3(−1G) differ only in one nucleotide, the −1 position of the P3 promoter, which leads to the same RNAIII transcript, just with different NAD modification levels. To test the influence of this single mutation on the transcriptome; RNA-Seq was performed on total RNA from both strains. Their transcriptomes were found to be very similar to each other, with only 4 upregulated genes (*splA*, *splB, splC, splD*, all of them coding for serine proteases, Table S8) and 2 downregulated genes (*spa,* an uncharacterized gene with homology to *ssaA,* Table S8) in *S. aureus* pCG-P3(−1G). The *splABCD* operon is known to be activated via RNAIII, whereas *spa* is known to be inhibited by RNAIII (Queck et al., 2008). Thus, the higher NADylation in pCG-P3(−1G) might enhance RNAIII’s inhibitory activity. Interestingly, the prototypic RNAIII target gene, *hla,* was neither upregulated nor downregulated in the RNA-Seq data. Hence, we speculated that NADylation of RNAIII may influence the translation of Hla, and therefore, an analysis of Hla abundance in culture supernatants was conducted. The Hla obtained in culture filtrates of *S. aureus* pCG-P3(−1G) was significantly lower than in *S. aureus* pCG-P3 in late exponential phase cultures (*t*-test; *P* = 0.0148, Figure 3C and Figure S1E). Thus, 5’-NAD capping of RNAIII might modulate the interaction between the 5’-UTRs of RNAIII and *hla* mRNA as well, leading to a lower translation of the latter RNA.

We next analysed whether NADylation impacts the translation efficiency of the RNAIII-encoded Hld. According to LC-MS of culture filtrates, half the amount of Hld was found in pCG-P3(−1G) (with the higher amount of 5’-NAD cap) compared to pCG-P3 (Figure 3C, Figure S1F). This finding indicates that RNAIII’s 5’-NAD cap might impair *hld* mRNA translation. A decreased production of two major hemolytic toxins, Hla and Hld, should lead to a less cytotoxic *S. aureus* strain. Indeed, an assay with *S. aureus* pCG-P3 and pCG-P3(−1G) culture supernatants in a human THP1 macrophage line (Figure 3D) revealed a significantly reduced cytotoxicity of pCG-P3(−1G) compared to pCG-P3 (Tukey’s multiple comparisons-test; *P* = 0.0004).

### 4.6 Hld’s Shine-Dalgarno sequence is accessible *in vitro* regardless of the presence of a 5’-NAD cap

In order to analyse whether 5’-NAD capping of RNAIII affects the secondary structure of its 5’-domain, we performed Selective 2’-Hydroxyl Acylation and Primer Extension (SHAPE) (Weeks & Mauger, 2011; Wilkinson, Merino, & Weeks, 2006). SHAPE is an approach to probe the structure of every nucleotide within an RNA simultaneously. The SHAPE reagent chemically modifies (acylates) the 2′-OH position of flexible nucleotides (Wilkinson et al., 2006). A subsequent DNA synthesis by reverse transcriptase stops one nucleotide before the position of a modified 2′-ribose position, reporting the site of a SHAPE modification in the RNA (Weeks & Mauger, 2011; Wilkinson et al., 2006). 1-methyl-7-nitro-isatoic anhydride (1M7) was chosen as SHAPE reagent (Mortimer & Weeks, 2007). To determine whether RNAIII’s secondary structure is modulated by the nature of its 5’-end, pure 5’-NAD-RNAIII and ppp-RNAIII were required. Pure full-length (514 nt) NAD-RNAIII could not be prepared, as no method exists for its preparative separation from full-length ppp-RNAIII. Therefore, we first probed a shorter version of RNAIII consisting of nucleotides 1 to 113, designated as RNAIII leader, which could be prepared by *in vitro* transcription and APB-gel purification as pure NAD-capped (NAD-RNAIII leader), or pure triphosphorylated version (ppp-RNAIII leader). To exclude the possibility that the truncated RNAIII leader sequences fold differently than the full-length RNAIII, we compared in a second step pure full-length ppp-RNAIII with impure full-length NAD-RNA (containing NAD-RNA and ppp-RNA ∼ 1:1). The SHAPE data showed very similar nucleotide reactivity profiles for NAD-RNAIII leader and ppp-RNAIII leader (Figure 4A, Figure 4B and Figure S1G), indicating an accessible SD sequence regardless of their 5’-end modification.

**Figure 4:**
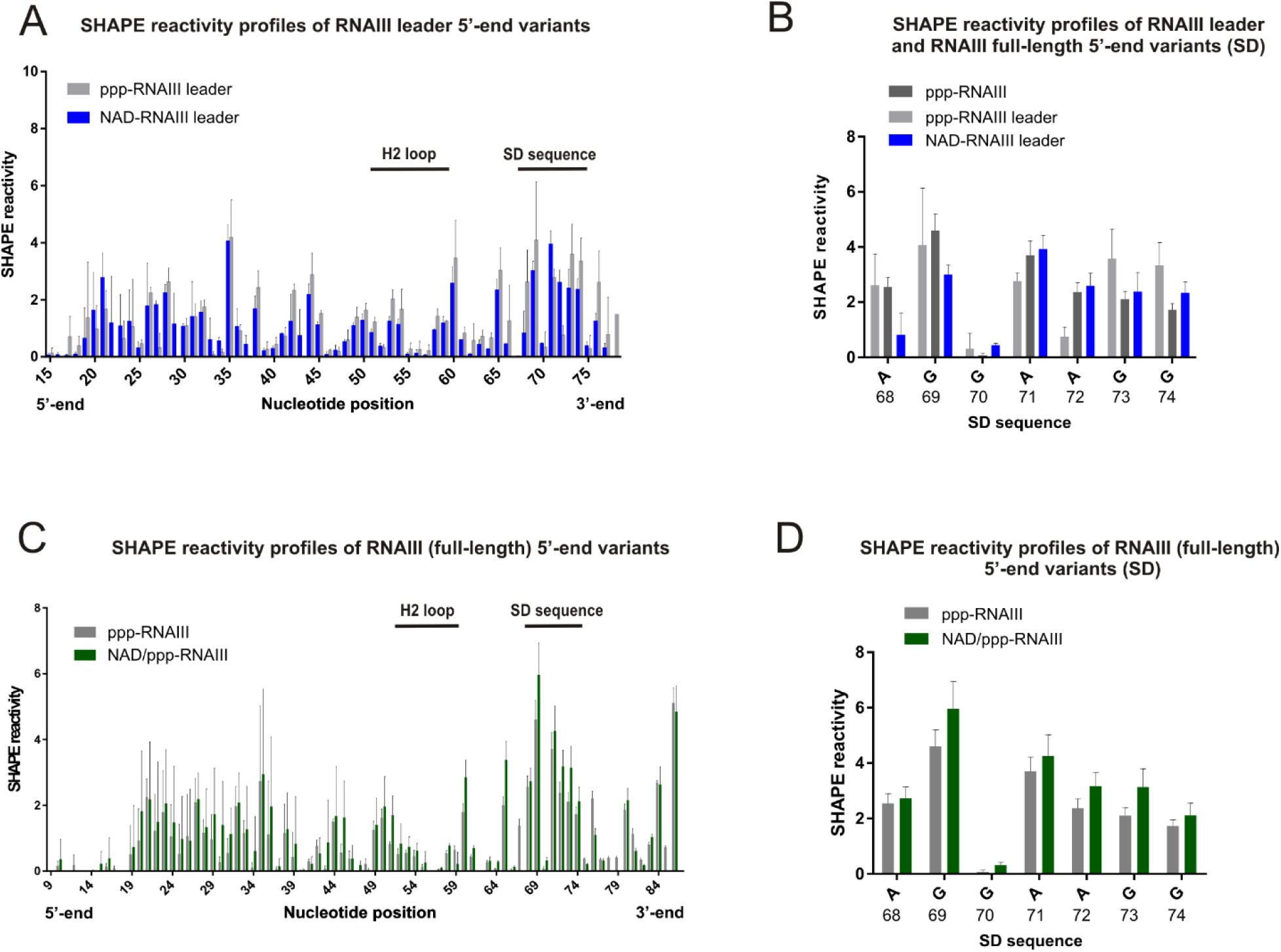
Analysis of the secondary structure of RNAIII 5’-end variants by SHAPE. **A:** The bar chart represents the differential SHAPE reactivity (Y-axis) throughout the nucleotides of the 5’-UTR of RNAIII (X-axis). The blue bars represent NAD-RNAIII leader whereas the light grey bars represent ppp-RNAIII leader (113 nt long). The region containing the SD sequence and the H2 loop residues are indicated as a horizontal black lines. **B:** Bar chart showing a detailed view of the SHAPE reactivity (accessibility) of the SD sequence from ppp-RNAIII leader (light grey bar) and NAD-RNAIII (blue bar). Full-length (514 nt) ppp-RNAIII is represented with dark grey bars. A: adenosine, G: guanosine. **C:** Same as panel A but the chart compares full-length ppp-RNAIII (grey bar) vs NAD/ppp-RNAIII (green bar) SHAPE experiments. **D:** Same as panel B but comparing the SD sequence accessibility of full-length ppp-RNAIII (grey bar) vs NAD/ppp-RNAIII (green bar). Horizontal bars represent mean. Error bars depict standard deviation.

When shaping full-length ppp-RNAIII and the ppp-/NAD-RNAIII mixture, both the overall SHAPE reactivity profile and SD’s accessibility were similar to that of the ppp-RNAIII leader (Figure 4C, Figure 4D)

## 5 Discussion

### 5.1 A short, regulatory, and protein-coding NAD-capped RNA in *S. aureus*

In this study, we have demonstrated the existence of 5’-NAD-capped RNAs in the opportunistic pathogen *S. aureus.* Since the discovery of the prokaryotic NAD cap in 2015, the presence of NAD-RNAs in many different organisms has been confirmed, and this modification might be ubiquitous. Nevertheless, the biological function of the 5’-NAD cap in prokaryotes is still unclear. In the case of *B. subtilis*, also a gram-positive bacterium, the NAD cap stabilises RNA against exoribonucleolytic attack by RNase J1 *in vitro* (Frindert et al., 2018). As in other organisms (Cahová et al., 2015; Frindert et al., 2018), several mRNAs were found to be enriched in the NAD captureSeq of *S. aureus* ATCC 25923. Most of these mRNAs encode proteins involved in different cellular processes: redox reactions, membrane transport, biosynthesis-related enzymes, phosphatases, kinases, and hydrolases (Table S2). Similar classes were observed in the *S. aureus* mutant strains (pCG-P3, pCG-P3(−1G) and pCG-P2), with several hits common with the wild type ATCC 25923 strain, i.e., NADP reductases, synthetases, and transcription factors.

Interestingly, the mutant-specific hits encoded for proteins that were similar to the ones found in the wild type strain, especially reductases such as the peptide methionine sulfoxide reductase or redox enzymes like the superoxide dismutase (Table S5, Table S6 and Table S7). Of particular interest is the enrichment of mRNAs that encode for NAD-NADP-utilizing enzymes (2-dehydropantoate 2-reductase, L-lactate dehydrogenase). A similar observation has been made in *B. subtilis,* where L-threonine 3-dehydrogenase (*tdh*) was enriched (Frindert et al., 2018). These findings underpin the possibility that some NAD-mRNAs might act as a cofactor of their encoded enzyme. Moreover, the NAD cap could act as an enzyme-binding site to its mRNA as an additional regulatory mechanism (Jaschke, Hofer, Nubel, & Frindert, 2016). However, the existence of this feedback mechanism still needs to be demonstrated empirically.

The *hld*/RNAIII gene was by far the most enriched of all hits. The results showed a higher NAD modification level of RNAIII in the wild type strain (∼36%) compared to pCG-P3 (∼10%). This phenomenon might be due to the different *S. aureus* strains used (ATCC 25923 and plasmid-carrying HG001) although these values were obtained by different approaches (LC-MS vs Northern blot). The variation in NAD content is supported by a different *hld/*RNAIII enrichment in NAD captureSeq between both strains after, with a higher value obtained in ATCC 25923 (Table S2, Table S5). Furthermore, a distinct NAD modification level on RNAIII could be the cause of some differences in the infective behaviour between the strains. Nevertheless, RNAIII is still strongly NAD-modified whenever transcribed by P3 promoter, and this together with its high expression level, its role as a crucial intracellular effector of the quorum-sensing system, and the additional presence of an embedded ORF for the *hld* gene (Figure 1A) make RNAIII a promising candidate for unveiling a biological function of the NAD cap.

### 5.2 The P3 promoter acts as a driver of NAD incorporation

The −1 position of the P3 promoter is found to modulate the incorporation of NAD into the nascent RNAIII. We also tested the effect of a promoter exchange on NAD transcriptional incorporation. The other promoter of the *agr* locus, P2, seems not to be prone to introduce NAD into RNAII (Log2FC = −1.65 in NAD captureSeq). These two promoters share several characteristics: phosphorylated AgrA activates transcription of both, they have a suboptimal interspace region between the -35 and −10 boxes, and both have been predicted to initiate with a +1A (Novick et al., 1995; Reynolds & Wigneshweraraj, 2011). Placing the P2 promoter upstream of the RNAIII sequence led to the total abolishment of NAD incorporation into RNAIII (Figure 2C and Figure 2E). The unexpected revelation of the TSS of the P2 promoter being G instead of A (Figure 2E) provided us with an additional “NAD zero” control in our experiments, constituting a transcript that differed only in one nucleotide from the native one but was devoid of any 5’-NAD. This control further demonstrated that NAD captureSeq is not biased towards highly expressed RNAs unless they bear the NAD cap. Moreover, it also confirmed the reliability of the APBgel-Northern blot combination as a tool for analysing NAD-capped RNAs.

### 5.3 NAD-RNAIII as a modulator of *S. aureus* physiology

For validating the function of RNAIII’s NAD cap, it was essential that the RNAIII expression of the different *S. aureus* strains was not altered by the changes introduced in the promoter sequence. qPCR analysis confirmed the lack of bias in RNAIII content between *S. aureus* pCG-P3 and pCG-P3(−1G) (Figure 3B). This experiment also showed lower RNAIII levels in the pCG-P2 strain (Figure 3B), which suggest that the P2 promoter is less active than the P3 promoter in these experiments.

This finding is in agreement with the fact that *in vitro,* AgrA-mediated activation of transcription is more prominent at P3 than at P2 (Reynolds & Wigneshweraraj, 2011). On the other hand, a recent *in vivo* study reported AgrA∼P affinity to P2 to be higher than to P3, previously assessed by Koenig et al. (2004), and this differential affinity being crucial for the *agr* positive feedback loop (Garcia-Betancur et al., 2017).

According to transcriptome sequencing, the higher NAD modification ratio of RNAIII in *S. aureus* pCG-P3(−1G) (Figure 2C and Figure 2E) appears to downregulate the transcription of *spa* (protein A) and *ssaA* (staphylococcal secretory antigen SsaA) (Table S6). *SsaA* mRNA has been proposed (but not yet demonstrated) to base-pair with the 3’-end of RNAIII (Lioliou et al., 2016). *spa* mRNA is known to base-pair with RNAIII, in particular with helix 13 (Huntzinger et al., 2005), thereby blocking *spa*’s Shine-Dalgarno sequence, leading to *spa* mRNA translation repression and degradation by endoribonuclease III (Bronesky et al., 2016). While within the primary sequence helix 13 is far away from the NAD-5’-end, in one of the proposed secondary structures of RNAIII, the 5’ and 3’-ends base-pair with each other (Novick et al., 1993), thereby localising the NAD in the vicinity of helix 13. Hence, it is conceivable that the 5’-NAD cap affects the folding of RNAIII and its interaction with the *spa* mRNA. A stronger interaction might lead to a decreased protein A production, which would ultimately favour the dissemination of *S. aureus* pCG-P3(−1G) isolates (Bronesky et al., 2016). Furthermore, four genes for a protease family occurring only in *S. aureus*, (*SplA/B/C/D* operon) (Reed et al., 2001) were upregulated in the strain with increased NAD-RNAIII. Also, the *Spl* operon is induced by the *agr-*system with a mechanism that has not been elucidated but that might involve RNAIII (Reed et al., 2001). The Spl proteases have been implicated in tissue dissemination processes due to their ability to degrade host proteins (Paharik et al., 2016). Noteworthy, a recent study reported a decrease in the amount of Hld in the spent medium obtained from an *S. aureus Spl* deletion mutant, suggesting a positive correlation between RNAIII and *SpI.* On the other hand, the *ΔSpl* strain did not show alterations in Hla levels, proving that the *agr* system was not inhibited (Paharik et al., 2016).

At this point, one could expect that the upregulation in Spl proteases in pCG-P3(−1G) together with the downregulation of protein A and SsaA would generate a more invasive bacterium than pCG-P3. Nevertheless, evidence at protein level of these two targets would be necessary to predict more accurately the phenotypic effects.

Transcriptome sequencing did not reveal significant changes in another major target of RNAIII, namely *hla* (Log2FC = -0.14). However, as RNAIII activates translation of *hla* mRNA by base pairing between the 5’-UTRs of both RNAs (Morfeldt et al., 1995), an effect on the mRNA level of this gene was not expected.

On the protein level, the phenotypic effects were more notorious. Since pCG-P3(−1G) exhibits an upregulated *Spl* operon, an increase in Hld amount in supernatant could be expected. In contrast, a significant decrease of Hld in cell culture filtrates of *S. aureus* pCG-P3(−1G) was observed (Figure 3C).

One explanation would be that the 5’-NAD cap in RNAIII might impair Hld translation. In eukaryotes, there are conflicting reports regarding the translatability of NAD-RNAs (Jiao et al., 2017; Wang et al., 2019), and there are no comparable data on prokaryotes which are not known to sense and require “cap” structures in translation. It has been suggested that the 3’-end of RNAIII blocks Hld translation by base-pairing with its Shine-Dalgarno (SD) sequence, making it inaccessible to ribosomes, thereby delaying the translation of Hld by 1h after RNAIII transcription (Balaban & Novick, 1995). Thus, the 5’-NAD cap might favour the interaction between the 5’-end and and some downstream sequences due to the pseudo-base-pairing of the nicotinamide moiety of the initiating NAD, which has been proposed to base-pair with C and T of DNA in the open promoter (Vvedenskaya et al., 2018). Hence, it appears plausible that the 5’-NAD cap of an RNA interacts similarly with C and uridine (U) residues inside the same molecule. Nevertheless, even if one assumes that NAD-RNA is not translated while ppp-RNA is, this could not explain how a 15% reduction in ppp-RNAIII in pCG-P3(−1G) compared to pCG-P3 (Figure 2C) can cause a 50% decrease in translation product yield. We assume that this effect may be due to yet unknown factors that modulate *hld* translation.

In the case of Hla, the changes observed in pCG-P3(−1G) compared to pCG-P3 are slightly lower than for Hld (Figure 3C), which might imply a lower influence of the 5’-NAD-cap on the RNAIII-*hla* mRNA interaction than on RNAIII translation. Indeed, the fact that the H2 hairpin of RNAIII, which is the one predicted to interact with *hla* mRNA, is only nine nucleotides upstream of *hld*’s SD sequence suggests that both might be similarly affected by the 5’-NAD cap. Like in the case of Hld translation, nicotinamide’s base-pairing flexibility might increase the probability of generating a 5’-end-3’-UTR duplex in RNAIII’s secondary structure, sequestering both the SD sequence and H2 helix, and explaining the phenotypes obtained in *S. aureus* pCG-P3(−1G) (Figure 3C and Figure 3D). Surprisingly, the *in vitro* analysis of RNAIII leader’s secondary structure by SHAPE did not reveal differences related to the 5’-NAD cap (Figure 4A). In both RNAIII leader variants, the SD appeared to be accessible (Figure 4B). This structure would likely be compatible with translation of the embedded Hld ORF regardless of the presence of the 5’-NAD cap. Of note, the analysis was done with a shortened version of RNAIII with the aim of detecting the presence of a mechanism that would block co-transcriptional translation. Furthermore, SHAPE measurements with full-length RNAIII revealed an accessible SD sequence as well (Figure 4B, Figure 4C, Figure 4D), an observation already reported in the chemical probing of RNAIII performed by Benito et al. (2000). Our data suggest that the lack of RNAIII’s 3’-domain would not exert any effect on Hld and Hla translation (Figure 4B, Figure S1G). In contrast, a mutant *S. aureus* strain with a truncated RNAIII devoid of its 3’-end domain (211 nt) showed an earlier Hld translation compared to the wild type strain (Balaban & Novick, 1995). Likewise, deletions within 3’-end region of RNAIII also inhibit Hla expression (Novick et al., 1993). Thus, although the *in vitro* RNAIII SHAPE experiments do not explain the obtained phenotypes with decreased Hld and Hla, in a regulated cellular co-transcriptional translation environment, the 5’-NAD cap could cause some differences and, induce different phenotypes.

While RNAIII has been studied intensively as a central regulator of bacterial physiology, a large number of different targets affected by it, and the diversity of mechanisms of action make it rather unlikely that only one stable secondary structure induces all these responses. Instead, inside the cell likely exists an equilibrium between different folds, each of them suited for specific purposes (Benito et al., 2000). The existence of such alternative folds has been hinted at by the inaccessibility of a fraction of *in vitro-transcribed* NAD-RNAIII to NudC cleavage, and its modulation by an unfolding / folding cycle (Figure 3A). Thus, the existence in the cell of an RNAIII subpopulation that is sensitive to its 5’-modification state and is responsible for both *hld* and *hla* activation remains plausible (Balaban & Novick, 1995).

The fact that the RNAIII sequence is highly conserved among *S. aureus* strains led us to hypothesize that the incorporation of NAD into RNAIII should be strictly regulated and therefore the evolution selected the −1T at P3 promoter. In such a way, *S. aureus* may ensure the right (intermediate) level of NAD incorporation into RNAIII and preserves its optimal ensemble of secondary structures. Assuming this, the 5’-NAD cap might serve as modulator of RNAIII’s secondary structure. Moreover, the NADylation of RNAIII is likely dependent on the bacterial redox state and might be a new mechanism to fine-tune *agr* activity.

### 5.4 Outlook

In this study, we have discovered another prokaryotic organism that possesses NAD-capped RNAs. However, it constitutes the first evidence of this phenomenon in pathogenic bacteria to date. More importantly, we have found that a crucial regulatory and protein-coding RNA (RNAIII) is strongly NAD-modified and that alterations on its NAD-modification state lead to significant effects that modulate *S. aureus’s* cytotoxicity that could be related to an alternative secondary structure of RNAIII. However, and despite these exciting findings, the role of 5’-NAD-RNAIII remains unclear. Further studies will be conducted to unravel how exactly the NAD cap changes RNAIII’s secondary structure and how this change contributes to the overall secondary structure ensemble of RNAIII. Furthermore, *in vivo* studies applying our mutant *S. aureus* strains to animal infection models are planned to study the pathogenicity of this bacterium.

## 6 Materials and Methods

### 6.1 Bacterial strains

*S. aureus* ATCC 25923 was the selected strain to detect the presence of NAD covalently linked to RNA by UPLC-MS analysis, NAD captureSeq, and pull-down of specific RNAs. *S. aureus* HG001 was used as wild-type and for the mutational studies. The pCG-246-based (Helle et al., 2011) shuttle plasmids pCG-P3, pCG-P2 and pCG-P3(−1G) were first grown in *E. coli* K-12 strain, then transformed into the restriction-deficient *S. aureus* RN4220, and finally electroporated into *S. aureus* HG001 *ΔRNAIII* as described before (Charpentier et al., 2004). For the cytotoxicity assay, the constructs were introduced into *S. aureus ΔRNAIII Δpsmαβ.* Genomic DNA from *S. aureus* ATCC 25923 was used to amplify the P3 promoter, P2 promoter, and RNAIII sequences (Q5 Hot Start High-Fidelity DNA Polymerase, NEB). The bacterial strains used in this study are summarised in Table S9.

### 6.2 Generation of constructs

The RNAIII sequence with the native P3 promoter sequence upstream was amplified from Genomic DNA from *S. aureus* ATCC 25923. The PCR amplification was carried out with the primers Fwd-P3-RNAIII/Rv-P3-RNAIII; as a consequence, the restriction sites *BamHI* and *EcoRI* were introduced in the amplicon. The P2 promoter sequence was fused to RNAIII by the following procedure: a PCR with the primers Fwd-P2/Rv-P2hyb was performed in order to generate a P2-amplicon with a 20 bp overhang on its 3’-end which was complementary to RNAIII’s first 20 nt. The PCR with primers Fwd-RNAIII/Rv-RNAIII originated a second amplicon containing the whole RNAIII sequence. Both amplicons were used as primers and subjected to PCR in order to generate the P2-RNAIII product, which contained *BamHI* and *EcoRI* restriction sites. Both PCR products were treated with *BamHI* and *EcoRI* and cloned into the pCG-246 shuttle vector (Helle et al., 2011) yielding the constructs pCG-671 and pCG-672. The RNAIII terminator sequence was introduced afterwards in both of the constructs by site-directed mutagenesis generating pCG-P3 and pCG-P2 plasmids. Besides, a mutation at the −1 position (−1T to −1G) respect to the TSS of the P3 promoter was inserted by site-directed mutagenesis, yielding plasmid pCG-P3(−1G). All primers used are summarised in Table S3.

### 6.3 NAD captureSeq and RNA-seq Next Generation Sequencing (NGS)

In order to identify those NAD-RNAs from *S. aureus*, we have made use of NAD captureSeq as described previously (Winz et al., 2017) and applied to 100 µg total RNA from *S. aureus* ATCC 25923 per sample isolated at the late-exponential phase (OD_600_ = 1.5). A NAD captureSeq variant was also used in order to compare the enrichment of specific RNAs between *S. aureus* strains (pCG-P2, pCG-P3, and pCG-P3(−1G)). For that, 15 fmol of variable length (61, 104, 205, 302, 400 nt) NAD-RNAs were spiked into each sample, acting as internal standards (IS). The PCR products were purified by polyacrylamide (PA) gel electrophoresis (PAGE), and the range between 150 and 300 bp selected (Figure S1A). This size selection helped to remove the primer dimers as well. Afterwards, the quality of the samples was analysed by Bioanalyzer measurements (Bioanalyzer 2100, Agilent Technologies, Santa Clara, California) using the Agilent DNA 1000 Kit (Agilent). The libraries were multiplexed and submitted to NGS on a NextSeq500 sequencer (Illumina, San Diego, California) with the following parameters: single-end reads, 75-bp read length, read depth 400 million. Besides, 20% of PhiX control DNA was spiked into the samples to provide sufficient read complexity. The NGS data analysis was done using an in-house pipeline. Briefly, after obtaining the raw reads, the leading guanosines were trimmed, and the 3’ adapter was removed by clipping. Once processed, the reads were mapped to the reference genome *Staphylococcus aureus subsp. aureus* NCTC 8325 (European Nucleotide Archive, Assembly: GCA_000013425.1) and to the IS RNA sequences (in the case of the quantitative NAD captureSeq, Table S4). The software used for mapping was Burrows-Wheeler Aligner (BWA-MEM, version: 0.7.13) (H. Li, 2013). To identify the gene hits, HTSeq (version: 0.6.0) (Anders, Pyl, & Huber, 2015) was used. Also, DESeq (Anders & Huber, 2010) was used for statistical analysis. Finally, the obtained hits were checked and visualised with the Integrated Genome Browser (Nicol et al., 2009). RNA-seq analysis was conducted with total RNA of *S. aureus* pCG-P2, pCG-P3, and pCG-P3(−1G) strains (biological triplicates, 5 μg total RNA each sample) isolated at the late-exponential phase (OD_600_ = 2.5). For removal of ribosomal RNA from samples, the Ribo-Zero rRNA Removal Kit (Gram-Positive Bacteria, Illumina) was used. The libraries were prepared with NEBNext Ultra II Directional RNA LibraryPrep Kit for Illumina (New England Biolabs (NEB), Ipswitch, Massachussets) together with the NEBNext Multiplex Oligos for Illumina (NEB). The removal of primer excess and primer dimers was done by the Agencourt AMPure XP RNA Clean beads (Beckman Coulter, Brea, California). The amount of sample was assessed by Qubit HS DNA assay (Thermo Fisher Scientific, Waltham, Massachussets), whereas the sample quality was checked with Agilent Tape Station D1000 for DNA. Samples were equimolarly pooled before 50 SE sequencing on the Illumina HiSeq platform. NGS data analysis was performed with an in-house pipeline, and the reads were mapped to the reference genome *Staphylococcus aureus subsp. aureus* NCTC 8325 (European Nucleotide Archive, Assembly: GCA_000013425.1).

### 6.4 Total RNA isolation

*S. aureus* strains were grown at 37 °C with shaking at 200 rpm in LB broth (Lennox), and cells were harvested at OD_600_ = 2.5 (experiments conducted with *S. aureus* HG001 plasmid-transformed strains) and OD_600_ = 1.5 (experiments conducted with *S. aureus* ATCC 25923). The RNA was extracted using TRIzol reagent (Thermo Fisher Scientific). Briefly, the pelleted cells were resuspended in TE buffer (30 mM Tris-HCl, 1 mM EDTA, pH = 8.0) supplemented with lysozyme 20 mg/mL (Sigma-Aldrich, Saint Louis, Missouri) and 80 μg/mL lysostaphin (Sigma-Aldrich) and incubated 30 min at 37 °C. The cells were incubated for 30 min at -80 °C before the addition of the TRIzol reagent. Afterwards, the protocol proceeded according to the manufacturer’s instructions. Samples were treated with DNase I (1.5 U/mg total RNA, Roche, Basel, Switzerland) at 37 °C for 1 hour to remove the genomic DNA, then they were P/C/I-chloroform-extracted and precipitated. The RNA was washed twice with 80% ethanol and dissolved in Millipore water. The quality of the RNA was assessed by agarose gel electrophoresis (Figure S1H). The concentration and purity of the total RNA were measured by Nanodrop (Thermo Fisher Scientific), paying attention to OD_260_/OD_280_ and OD_260_/OD_230_ ratios.

### 6.5 Quantitative PCR

To confirm the enrichment of the obtained hits at the cDNA level and thereby rule out the possible PCR bias, qPCR measurements were carried out as described before (Cahová et al., 2015). 20 µL reactions were performed in triplicate using 3 µL cDNA (1:50 diluted) as a template. When confirming the cDNA enrichment of the hits, APDRC-treated sample cDNA with cDNA of the minus ADPRC control were compared by the 2^-ΔΔC^_T_-method (Livak & Schmittgen, 2001). The 5SrRNA gene was used as internal control and Millipore water as a negative control. Another qPCR assay was performed to check the *hld*/RNAIII levels of the different *S. aureus* strains (pCG-P2, pCG-P3, and pCG-P3(−1G)). 5 μg of total RNA (harvested at late-exponential phase, biological triplicates) from each strain was reverse transcribed with Superscript IV reverse transcriptase (Thermo Fisher Scientific) following the manufacturer’s instructions. 3 µL cDNA (1:100 diluted) per 20 µL reaction were used as the template. The *gyrB* gene was used as internal standard and Millipore water as a negative control. Standard curves were determined for each gene, using purified chromosomal DNA at concentrations of 0.005 to 50 ng/ml. All qPCR experiments were performed in a Light Cycler 480 instrument (Roche) using the Brilliant III Ultra-Fast SYBRGreen qPCR Mastermix (Agilent Technologies). The data were analyzed with the Light Cycler 480 Software (Agilent Technologies). The primers used for qPCR analysis are listed in Table S3.

### 6.6 Detection of Hld by HPLC/ESI-MS

The detection of Hld levels from different *S. aureus* strains was performed by high-performance liquid chromatography/electrospray ionisation mass spectrometry (HPLC/ESI-MS) of overnight culture filtrates as described earlier (Queck et al., 2008) with minor changes. The column (Zorbax SBC8, 2.1 x 50 mm, 3.5 μ, Agilent Technologies) was run at 0.3 ml/min with a gradient of 0.1% trifluoroacetic acid in water to 0.1% trifluoroacetic acid in acetonitrile. HPLC-MS experiments were performed on a Bruker microTOFQ-II ESI mass spectrometer (Bruker, Billerica, Massachussets) connected to an Agilent 1200 Series HPLC system equipped with a multi-wavelength detector (Agilent Technologies). ESI was used with a capillary voltage of 4,500 V, and the collision voltage was set to 10 eV. The drying gas temperature was 200 °C, the drying gas flow rate was 6 Lmin^-1,^ and the detector was operated in positive ion mode. Mass spectra were recorded on a range from 250 m/z to 3,000 m/z. The two m/z peaks in the mass spectrum for Hld (formylated and deformylated version, Figure S1F) were used to calculate PSM concentration by integration using ACD/Spectrus software (Advanced Chemistry Development (ACD), Toronto, Canada).

### 6.7 Western blotting of Hla

One milliliter of cell culture filtrates from each strain (harvested at OD_600_ = 2.5) was filtrated and concentrated (final volume 20 μL) through ultracentrifugal filters (30 kDa MWCO, Amicon, Merck, Darmstadt, Germany) and analysed by 10% SDS-PAGE. After the run, SDS-PA gels were blotted onto Amersham Hybond P 0.2 μm PVDF membranes (GE Healthcare, Chicago, Illinois) using an EasyPhor Semi-Dry-Blotter (Biozym Scientific, Hessisch Oldendorf, Germany). Membranes were blocked in TBST (Tris-buffered saline with 0.1% Tween 20) supplemented with 5% milk powder (Carl Roth, Karlsruhe, Germany). Afterwards, membranes were incubated with anti-staphylococcal alpha-toxin rabbit antibody (1:5000, Sigma-Aldrich) in washing buffer (TBST with 1% milk powder). After washing, Alexa Fluor Plus 488-conjugated goat anti-rabbit IgG (Thermo Fisher Scientific) in washing buffer (1:10000) was applied over the membranes. Membranes were washed with 50 mM Tris-HCl, pH 7.25, and subsequently scanned using a Typhoon FLA 9500 imager (GE Healthcare). Quantification was done with ImageQuant TL software (GE Healthcare).

### 6.8 Detection of NAD covalently bound to RNA by UPLC/MS

Samples (total RNA and pull-down RNA) were prepared as described before (Frindert et al., 2018). Briefly, RNA was extensively washed with decreasing urea concentrations in ultracentrifugal filters (10 kDa MWCO, Amicon, Merck). RNA was recovered and subjected to NudC and alkaline phosphatase (Sigma-Aldrich) treatment (1 h at 37 °C, 2 h for pull-down RNA) in the presence of the MS internal standard (d4-riboside nicotinamide, Toronto Research Chemicals, Ontario, Canada). Afterwards, the reaction mixtures were filtered through 10 kDa ultracentrifugal filters (Merck) and dried under reduced pressure. Then, samples were analysed by UPLC-MS following the previously described protocol (Frindert et al., 2018). A calibration curve was recorded for r-NA for each analytical batch. M/z peaks of the mass spectra from the analyte and internal standard were integrated with TargetLynx software (Waters corporation, Milford, Massachussets). The amount of injected r-NA was calculated by integration of the corresponding m/z peak using the TargetLynx software and the calibration curve.

### 6.9 Gel electrophoresis and Northern Blot analysis

RNA was separated by denaturing PAGE. For NAD captureSeq NGS library generation (Qualitative and Quantitative), the cDNA amplification products were purified by native PAGE. APBgels (Nübel et al., 2017) were used to separate and purify NAD-RNA and ppp/p-RNA. Northern blot analysis was performed as described before (Cahová et al., 2015). RNA was separated either by PAGE or PAGE-APBgels (200 V, 90 min), blotted onto a Whatman Nytran SuPerCharge nylon blotting membrane (Merck) for 3 h (4 h for APBgels at 4 °C to avoid buffer evaporation) at 250 mA using an EasyPhor Semi-Dry-Blotter (Biozym Scientific), and UV-crosslinked. Membranes were pre-hybridized in Roti-Hybri-QuickBuffer (Carl Roth) for 2 h at 48 °C. Afterward, 5 µL RNA radiolabeled probe (Table S3) was added and incubated overnight at 48 °C in a hybridization oven with rotation. The templates for the probes were prepared by PCR using Taq DNA polymerase (prepared laboratory stock). Probes were prepared by *in vitro* transcription (IVT) in the presence of 35 μCi α-^32^P-ATP and 35 μCi α-^32^P-CTP (3,000 Ci/mmol, each, Hartmann Analytic, Braunschweig, Germany). The IVT reactions were treated with DNase I (Roche), P/C/I-chloroform-extracted, ethanol-precipitated and dissolved in 30 µL Millipore water. The blot was washed for 30 min with wash solution 1 (2x SSC, 0.1% SDS), 30 min with wash solution 2 (1x SSC, 0.1% SDS) for 30 min with wash solution 3 (0.25x SSC, 0.1% SDS). Radioactive RNA was visualized using storage phosphor screens (GE Healthcare) and a Typhoon FLA 9500 imager (GE Healthcare).

### 6.10 Preparation of NAD-RNAIII and ppp-RNAIII markers and IS NAD-RNAs

The RNAIII markers used in the Northern blot analysis were prepared by IVT. The IVTs were performed as described earlier (Huang, 2003). Each 100 μl IVT reaction contained: 1.5-2 µg DNA template and 4 mM of each NTP (6 mM NAD for NAD-RNAs; ATP concentration was reduced to 2 mM, or without NAD for ppp-RNAs). The reactions were performed in transcription buffer (40 mM Tris-HCl pH = 8.1, 1 mM spermidine, 22 mM MgCl_2_, 0.01% Triton X-100, 10 mM dithiothreitol (DTT), 5% DMSO). Reactions were stopped upon addition of denaturing gel loading buffer (10% TBE in formamide containing 0.05% bromophenol blue, 0.5% xylene cyanol blue) and purified by PAGE with standard running conditions (1x TBE buffer, 600 V). RNAs were excised after the run, eluted overnight in 0.3 M sodium acetate (pH = 5.5) at 19 °C, isopropanol-precipitated and dissolved in Millipore water. DNA templates for the RNAIII markers were prepared by PCR amplification of genomic DNA from *S. aureus* ATCC 25923 using Q5 Hot Start High-Fidelity DNA Polymerase (NEB) and the primers listed on Table S3. The PCR products were purified with the QIAquick PCR purification kit (Qiagen, Hilden, Germany) before the IVT reactions.

The 5 different IS NAD-RNAs used for NAD captureSeq were also prepared by IVT (Huang, 2003). As a template, regions of *E. coli’s* K12 genome with no homology within *S. aureus* genome were PCR amplified (Q5 Hot Start High-Fidelity DNA Polymerase (NEB)) with the primers summarised in Table S3. The PCR products were purified with QIAquick PCR purification kit (Qiagen) before IVT. IVT reactions were PAGE purified, as described above. Afterwards, the RNAs were additionally purified on APBgels (5% PA, 0.4% APB) (Nübel et al., 2017), loaded with APBgels loading buffer (8 M urea, 10 mM Tris-HCl pH = 8.0, 50 mM EDTA, xylene cyanol and bromophenol) and run in 1x TAE (550 V). The sequences of each IS NAD-RNA are summarised in Table S4.

### 6.11 Preparation of radiolabeled pure NAD-RNAIII and NudC cleavage assay

The radiolabeled (body labelled) pure NAD-RNAIII was prepared by IVT essentially as described before (Huang, 2003) but with the presence of 100 μCi α-^32^P-UTP (3,000 Ci/mmol, Hartmann Analytic) per 100 μL reaction. The IVT reaction was first PAGE-purified and afterwards purified on APB gels as described above to obtain 100% NAD-modified RNAIII, which was subjected to NudC cleavage. The NudC cleavage assay was carried out as described before (Cahová et al., 2015) but with a ratio of 2 NudC:1 RNAIII. 10 μL reactions per triplicate in 1x degradation buffer (25 mM Tris-HCl pH = 7.5, 50 mM NaCl, 50 mM KCl, 10 mM MgCl_2_, 10 mM DTT) were incubated for 5, 10, 20, 30, 60 and 120 min at 37 °C. Reactions were stopped by addition of APBgel loading buffer and analysed on APBgels (6% PA, 0.7% APB). The stability of NAD-RNAIII in 1x degradation buffer was assessed by incubating it for 2 hours at 37 °C in the absence and the presence of the enzyme (Figure 1, Figure 3A).

Additional reactions per triplicate were performed to test the influence of RNAIII’s secondary structure on NudC cleavage capacity. After a 60 incubation at 37 °C in the presence of NudC, samples were heated up to 75 °C for 2 min and cooled down to 25 °C (NAD-RNAIII folding after digestion). Afterward, fresh NudC was added, and samples underwent a second round of incubation (60 min at 37 °C, Figure 1, Figure 3A).

Radioactive RNA was visualised using storage phosphor screens (GE Healthcare) and a Typhoon FLA 9500 imager (GE Healthcare).

### 6.12 Pull-down of RNAIII and 5SrRNA

RNAs were specifically isolated out of 300 μg total RNA as described previously (Cahová et al., 2015; Frindert et al., 2018). Briefly, 150 μL Streptavidin Sepharose High Performance beads (GE Healthcare) were loaded onto Mobicol Classic columns (MoBiTec, Göttingen, Germany), washed with 1x PBS and supplemented with biotinylated DNA probes (Biomers, Ulm, Germany, Table S3) dissolved in 1x PBS (75 μL, 25 mM, with incubation for 10 min at 25 °C with shaking at 1000 rpm). After washing and equilibration of the beads (with 1x PBS and pull-down buffer (10 mM Tris-HCl pH 7.8, 0.9 M tetramethylammoniumchloride, 0.1 M EDTA pH 8.0), total RNA dissolved in pull-down buffer was added. RNA was then incubated for 10 min at 65 °C and then for 25 min at room temperature with rotation (Tube Rotator, VWR, Radnor, Pennsylvania). Beads were washed with Millipore water, and RNA was eluted upon addition of 2mM EDTA (pre-heated to 75 °C). The eluted RNA was precipitated with 0.5 M ammonium acetate and isopropanol in a reaction tube and dissolved in Millipore water. Northern blot analysis with the pull-down eluates and probes vs RNAIII and 5SrRNA was performed (Figure S1I). Afterwards, RNA was further purified by 8% PAGE and the bands corresponding to RNAIII and 5SrRNA excised (Figure S1J). After RNA precipitation and washing, RNA concentration was measured with Qubit RNA BR Assay Kit (Thermo Fisher Scientific). The purified RNA was used for: Northern bot analysis (Figure S1K) and NudC treatment with subsequent UPLC-MS analysis as described above. For DNAzyme treatment, the precipitated RNA was used without further PAGE purification (Figure 1E).

### 6.13 Generation of RNA-cleaving DNA enzymes

Four different RNA-cleaving DNAzymes were generated following the instructions by Joyce (Joyce, 2001) to generate a shorter RNAIII. DNAzymes were tested on RNAIII *in vitro,* and the most efficient one was selected for further experiments (DNAzyme II, Figure S1L). The cleavage of RNAIII by DNAzyme II (37 nt, Table S3) generated a 125 nt product containing RNAIII’s 5’-terminus. For the *in vitro* cleavage assays, 10 μL reactions were set up in 1x DNAzyme buffer (10 mM Tris-HCl pH = 8.0, 5 mM NaCl) supplemented with 1 μM NAD/ppp-RNAIII, 0.2 mM DTT, 25 mM MgCl_2_, and 1 μM DNAzyme II. First, RNAIII was folded in the presence of 0.2 mM DTT by incubation for 2 min at 75 °C, followed by a cool down to 25 °C. Afterwards, the remaining reagents were added (MgCl_2_, DNAzyme II) and reactions were incubated 1 h at 37 °C. Reactions were stopped by addition of denaturing gel loading buffer and analysed by classical PAGE and APBgels followed by Northern blot as described above (Figure S1L). The same procedure was followed with RNAIII obtained after pull-down (biological triplicates, 2 μM DNAzyme II) but including an additional NudC treatment (1.5 µM NudC, 1h 37 °C) right after RNA cleavage as a negative control (to deplete NAD-RNAIII). NAD-RNAIII and ppp-RNAIII treated with DNAzyme II and I were used as reference markers in the Northern Blot assay of RNAIII pull down samples.

### 6.14 Analysis of 5’-ends by circular RT-PCR and Sanger sequencing

To analyse RNAIII’s TSS on the different *S. aureus* mutant strains (pCG-P3, pCG-P2, pCG-P3(−1G)), circular RT-PCR was carried out as described before (Slomovic & Schuster, 2013) with small differences. In essence, 10 μg of total RNA of each strain was treated with RNA 5’-Polyphosphatase (Epicentre, Madison, Wisconsin). The dephosphorylated RNAs were self-ligated by T4 RNA ligase (Thermo Fisher Scientific) treatment, and the predicted 3’-end-5’-end ligated region of RNAIII was amplified by nested PCR with the primers listed in Table S3. The purified amplicons were then phosphorylated by T4-Polynucleotide Kinase (Thermo Fisher Scientific) treatment before cloning them into the plasmid pDisplay-AP-CFP-TM (Addgene plasmid #20861 (pDisplay-AP-CFP-TM was a gift from Alice Ting (Howarth, Takao, Hayashi, & Ting, 2005)), previously digested with EcoRV (NEB) and dephosphorylated by alkaline phosphatase (Thermo Fisher Scientific)). The blunt-end ligation was carried out by T4-DNA ligase (Thermo Fisher Scientific). The ligation mixture was transformed into *E. coli* DH5_α_ competent cells. The transformed cells were streaked on Petri dishes containing LB-agar supplemented with ampicillin (100 μg/mL, Carl Roth) and incubated overnight at 37 °C. Single colonies were picked, the plasmids isolated and subjected to high throughput Sanger sequencing (Microsynth SeqLab, Göttingen, Germany). All the enzymatic treatments were performed according to the manufacturer’s instructions.

### 6.15 Cytotoxicity assay

Overnight cultures were sub-cultured to OD_600_ = 0.05 in LB-broth (Lennox) and then grown for ∼ 22 h. Bacterial supernatants were harvested and filtered through a 0.45 µm filter (Merck) and analysed for its cytotoxic potential. The cytotoxicity assay was performed essentially as described (Munzenmayer et al., 2016) with the exception that 1×10^5^ THP1 cells were seeded in 96-well cell 8culture plates in a final volume of 100 µL. Differentiated THP1 macrophages were treated with 200 µL of each bacterial supernatant. THP1 cells with culture medium were used as negative control and THP1 cells treated with Phosphate Buffered Saline (PBS) containing 1% TritonX-100 were used as a positive control. The cytotoxic potential of the samples was determined from THP1-cell supernatants after 3 hours by using the Cytotoxicity Detection Kit (Roche) according to the manufacturer’s instructions.

### 6.16 Structural analysis of RNAIII variants by SHAPE

1M7 was synthesized as described (Mortimer & Weeks, 2007). A shorter version of RNAIII comprising nucleotides 1 to 113 (RNAIII leader) and full-length RNAIII with different 5’-ends (5’-NAD cap or triphosphate) were prepared by *in vitro* transcription as described above. In the case of full-length RNAIII, a mixture containing ∼ 1:1 NAD-RNAIII and ppp-RNAIII was prepared whereas the obtained NAD-RNAIII leader was 100% NAD-modified. The IVT template was generated by PCR with the primers RNAIII-leader-FW and RNAIII-leader-RV. Genomic DNA from *aureus* ATCC 25923 was used for the PCR. The NAD-supplemented IVT reaction was double-purified, first by denaturing PAGE and afterwards by APBgels, yielding NAD-RNAIII leader and ppp-RNAIII leader (Figure S1M).

SHAPE was performed as described with minor modifications (Wilkinson et al., 2006). Briefly, 8 pmol of RNA dissolved in 47.8 μL Millipore water were incubated for 2 min at 75 °C and then cooled down to 60 °C with a ramp of 0.1 °C/s. Afterwards, 24.2 μL of Dulbecco’s Phosphate Buffered Saline (DPBS, Sigma-Aldrich) supplemented with 3 mM MgCl_2_ were added to the samples, and they were incubated 2 min at 60 °C. Samples were then cooled down to 30 °C (0.1°C/s) and kept 2 min at this temperature. During this step, each sample was equally divided into two vials (sample vial and control vial). The vials were heated up to 37 °C, and then 4 μL of 100 mM 1M7 (10 mM final concentration) was added to sample vial. 1M7 was dissolved in dry DMSO (Sigma-Aldrich), so the reaction control was supplemented with the same volume of dry DMSO as in the sample (4 μL). The reaction mixtures were incubated for 20 min at 37 °C. Finally, reactions were stopped by adding 1 reaction volume of Millipore water, and the RNA was precipitated by ethanol in the presence of 0.3 M sodium acetate. The precipitated RNA was washed twice with 80% ethanol and dissolved in Millipore water to a concentration of 0.5 pmol/mL.

For primer extension, 100 pmol of SHAPE-RT1 primer (Table S3) were radioactively labeled on its 5’-end with 3 µL of γ-^32^P-ATP (3,000 Ci/mmol, Hartmann Analytic) per 20 μL reaction by using T4 polynucleotide kinase (Thermo Fisher Scientific), and further purified with the QIAquick Nucleotide Removal Kit (Qiagen). Both steps were performed following the manufacturer’s instructions.

The primer extension assay was applied to the RNA samples by using Superscript IV reverse transcriptase (Thermo Fisher Scientific). The reaction volume was set to 10 μL. 1 pmol of treated RNA per reaction was used as a template for the reaction together with 5 μL of 5’-end radiolabeled DNA primer 1 μM. A sequencing ladder was generated by reverse transcription of 1 pmol non-treated *in vitro-*transcribed ppp-RNAIII leader supplemented with 0.5 mM deoxynucleotide triphosphate mix each (Sigma-Aldrich), and 1 mM of the corresponding dideoxynucleotide triphosphate (Jena Bioscience, Jena, Germany). Reactions were performed according to the manufacturer’s instructions. The removal of residual RNA was done by addition of 2 μL 1 M NaOH to the samples followed by a 5 min incubation at 90 °C. Samples were then neutralised by the addition of 2 μL 1 M HCL and stored at -20 °C in denaturing gel loading buffer before being analysed by PAGE.

Samples processed in triplicate were analysed by 15% PA sequencing gels with standard running conditions (TBE buffer, 2000 V, 4h 30 min run time, Figure S1L). Radioactive cDNA was visualised with storage phosphor screens (GE Healthcare) and a Typhoon FLA 9500 imager (GE Healthcare). Quantification of PA sequencing gels was done with the software SAFA (Laederach et al., 2008). The intensities of the bands were normalised, and the control lanes (DMSO treated) were subtracted from the sample lanes (1M7 treated) to obtain the SHAPE values for each nucleotide. The SHAPE values for each nucleotide were afterwards used in the RNAstructure software (Reuter & Mathews, 2010) to create a structure prediction for the RNA sequence.

### 6.17 Statistical analysis

Except for NGS data analysis, where DESeq statistics was used (Anders & Huber, 2010), all statistics were analysed using the software Prism 6 version 6.01 (GraphPad, San Diego, California). Error bars depict standard deviations in all experiments. For specific data sets, identification of outliers was performed by Grubbs’ method (Alpha = 0.2). When comparing two groups, the parametric unpaired two-tailed Student’s t-test was selected. For the analysis of experiments involving three or more groups, the parametric one-way ANOVA test was done. The applied *Post hoc* analysis was Tukey’s multiple comparisons-test. Differences were considered significant when *P* < 0.05.

## 7 Author contributions

H.M. and A.J. designed the study. H.M., N.K., Y.Z., and G.N. performed the experiments. H.M., Y.Z., G.N., S.E.G., C.W., and A.J. analyzed the data. H.M. and A.J. wrote the initial draft of the manuscript, and H.M., A.J., C.W and S.E.G. edited the manuscript.

## 8 Acknowledgements

This work was financed by Baden-Württemberg Stiftung (grant BWST_NCRNA_045). We thank all the members of the Jäschke Group for their support and input during the discussions, A. Dalpke and K. Kubatzky for the BSL2 laboratory workspace, the CellNetworks Deep Sequencing Core facility, in particular D. Ibberson, for cDNA library preparation and sequencing, M. Brunner for the access to the LightCycler and S. Suhm for helping with the synthesis of 1M7 and APB. H.G.M.F. thanks the Deutscher Akademischer Austausch Dienst (DAAD) for the awarded scholarship.

---

**Figure S1:**
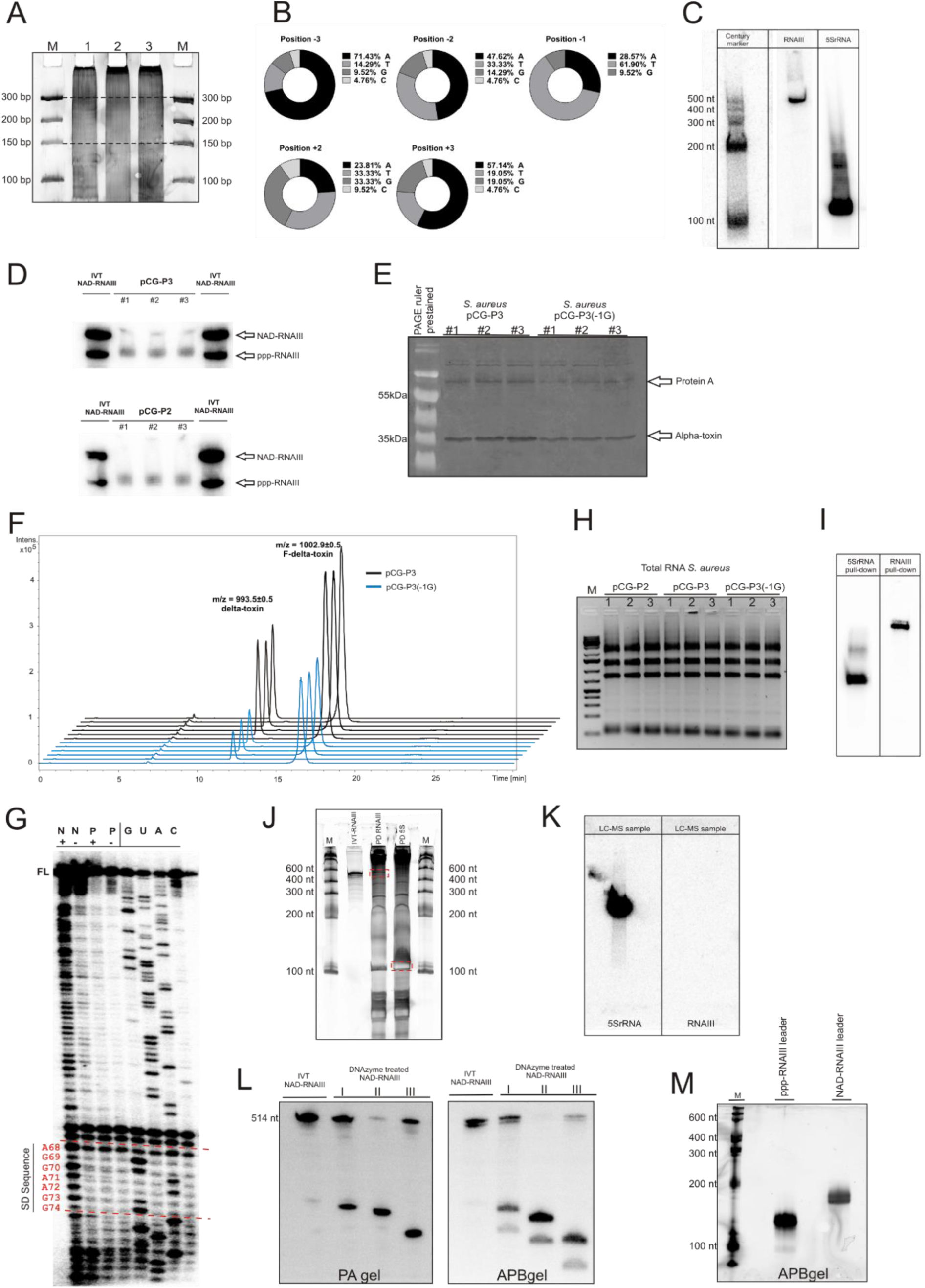
**A:** Size selection step during purification of PCR-amplified libraries with NGS primers on a 10% non-denaturing PA gel stained with SYBR Gold (Thermo Fisher Scientific). The dashed lines indicate the excised area (between 150 and 300 bp). The numbers indicate three replicates of minus ADPRC controls. M: Ultra Low Range DNA Ladder (Thermo Fisher Scientific). **B:** TSS analysis of the 21 hits enriched in NAD captureSeq in *S. aureus* ATCC 25923. Note that position +1 is not shown because all hits were selected to start with adenosine. A: adenosine, T: thymidine, B: guanosine, C: cytidine. **C:** Northern blot with probes against RNAIII and 5SrRNA. Ten micrograms of total RNA from *S. aureus* ATCC 25923 were loaded on each of the probe-hybridised lanes. Radioactively labeled RNA Century Marker (Thermo Fisher Scientific) was loaded on the first lane by the left. **D:** APBgels-Northern blot conducted in triplicate with total RNA of *S. aureus* pCG-P3 and pCG-P2 strains and probe against RNAIII. IVT Full-length mixed ppp/NAD-RNAIII was loaded as a control in the flanking lanes. Note that this experiment shows full-length RNAIII with no DNAzyme treatment. #1, #2 and #3 represent biological replicates. **E:** Membrane used for the detection of alpha-toxin (Hla) by Western blotting in culture filtrates from two strains (pCG-P3 and pCG-P3(−1G)) in triplicate (#1, #2 and #3). The marker used was the PageRuler Prestained Protein Ladder (Thermo Fisher Scientific). Two arrows indicate the bands corresponding to Hla (target) and protein A (unspecific binding of the antibody). **F:** Stacked extracted ion chromatograms from two different strains in triplicates (blue line pCG-P3(−1G), black line pCG-P3). The first peak corresponds to delta-toxin (Hld) whereas the second belongs to its formylated version. **G:** Analysis of SHAPE-teated RNAIII leader (*in vitro*). G, U, A and C represent the RNA nucleobases, which are equivalent to their cognate dideoxynucleotide used for the ladder generation. The bases of the SD sequence are highlighted in red. N: NAD-RNAIII leader treated with 1M7 (+) or with DMSO (-), P: ppp-RNAIII leader treated with 1M7 (+) or with DMSO (-), FL: Full length reverse transcription product (113 nt). **H:** Agarose gel electrophoresis analysis of total RNA samples from the different *S. aureus* strains Total RNA was isolated and treated with DNaseI as described in the methods section. M: 1 Kb Plus DNA Ladder (Thermo Fisher Scientific). Numbers indicate three replicates of each strain. **I:** Northern Blot of RNA pull-down eluates before PAGE purification. Samples were hybridised with RNAIII and 5SrRNA radiolabeled probes. **J:** The figure shows an 8% denaturing polyacrylamide gel with the pull-down products of RNAIII (PD RNAIII) and 5SrRNA (PD 5S). As a size marker ppp-RNAII, I was loaded, together with Riboruler Low Range RNA Ladder (Thermo Fisher Scientific). The red dashed areas indicate the excised region of the gels, corresponding to RNAIII and 5SrRNA. **K:** LC/MS-ready (after PAGE purification and band excision) RNAIII and 5SrRNA were subjected to Northern Blot detection. RNAIII concentration was too low to be detected. **L:** Screening of different DNAzymes (I, II, and III) on mixed ppp/NAD-RNAIII. The figure shows a Northern blot targeting RNAIII (probe complementary to first 86 nt of RNAIII), after electrophoresis on 8% polyacrylamide gel (left side) and APBgel (right side). The same ppp/NAD-RNAIII was loaded without treatment in both gels as a marker. The APB gel shows the separation between NAD and ppp RNAIII 5’-fragments. DNAzyme II is the most active of the three, acting quantitatively. **M:** Analysis of purified NAD / ppp-RNAIII leader on an APBgel. The ladder used was Riboruler Low Range RNA Ladder (Thermo Fisher Scientific).

**Table S1:** Detection of NAD by UPLC/MS/MS in total RNA samples from *S. aureus*. I and II: reactions by duplicate performed with NudC. Column 2 shows the NudC-treated samples minus the negative control (non-treated). The third column shows the estimated NAD-RNA molecules per cell.

| | Sample (fmol NAD/μg RNA) | Sample - negative control (fmol NAD/μg RNA) | Estimated NAD-RNA molecules per cell |
| --- | --- | --- | --- |
| *S. aureus* I | 31.52 | 23.61 | 839 |
| *S. aureus* II | 34.81 | 26.89 | 955 |

**Table S2:**
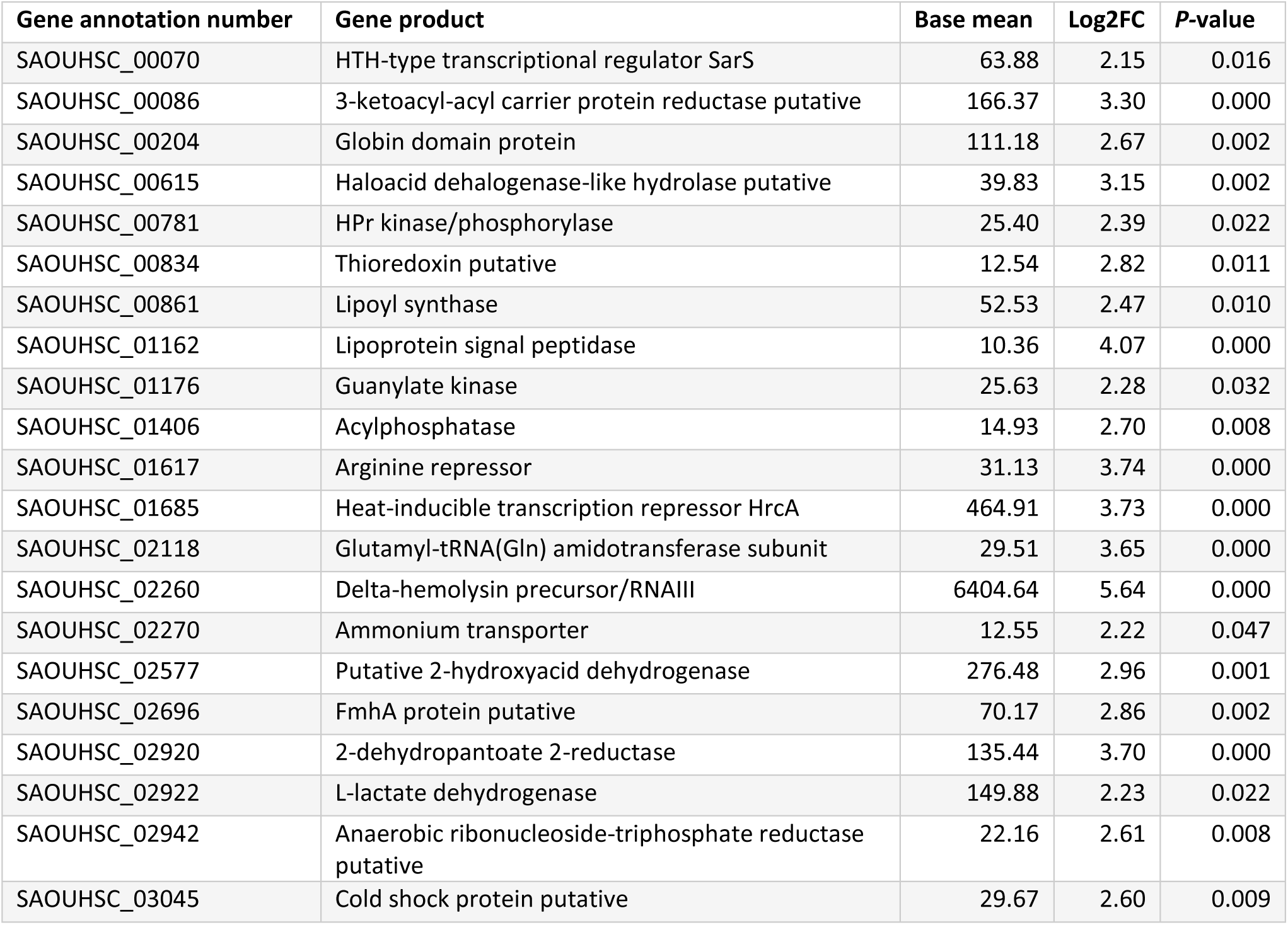
Enriched hits after NAD captureSeq in *S. aureus* ATCC 25923 strain. Base mean: normalised mapped reads, Log2FC: Log2 Fold change (sample vs negative control).

**Table S3:**
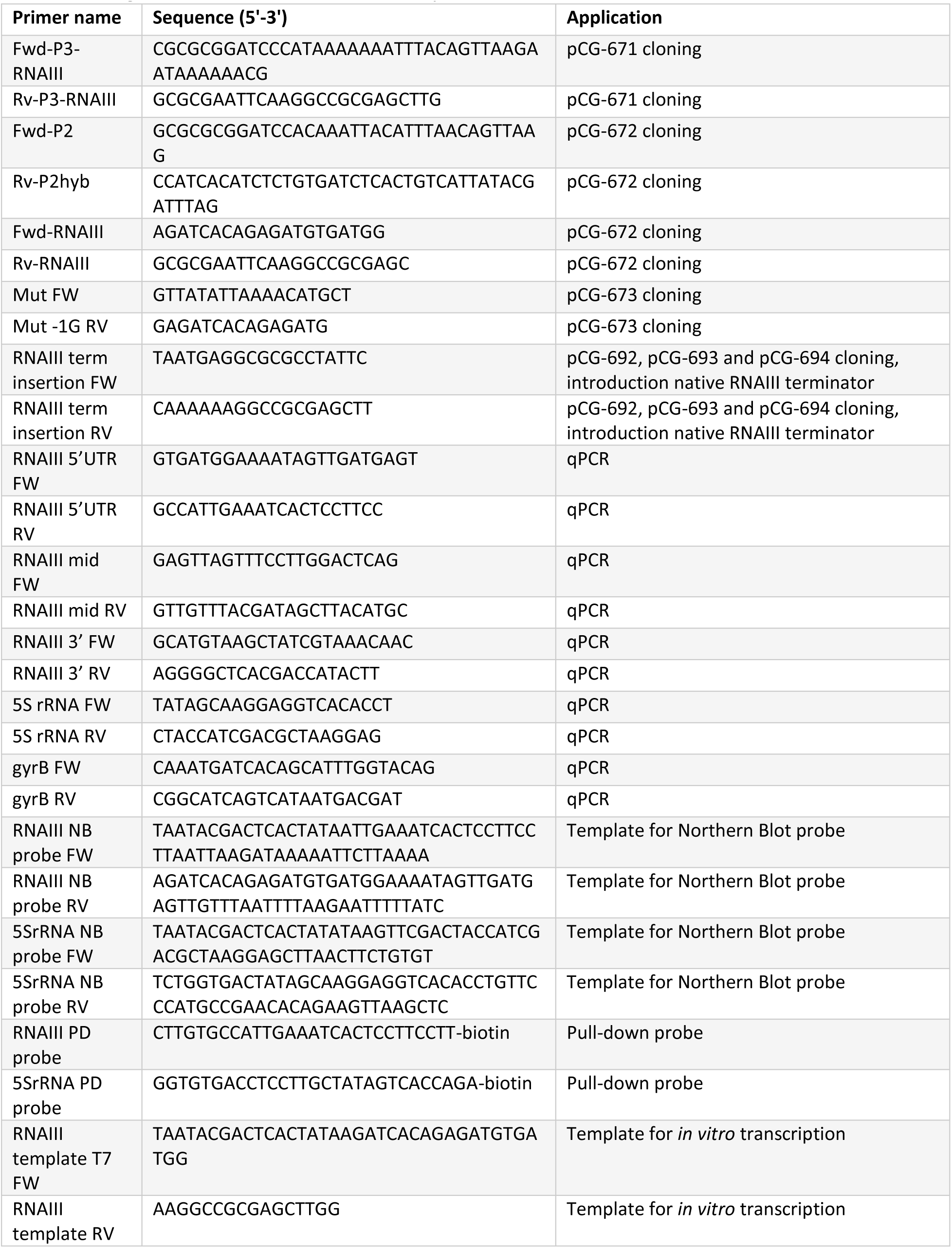
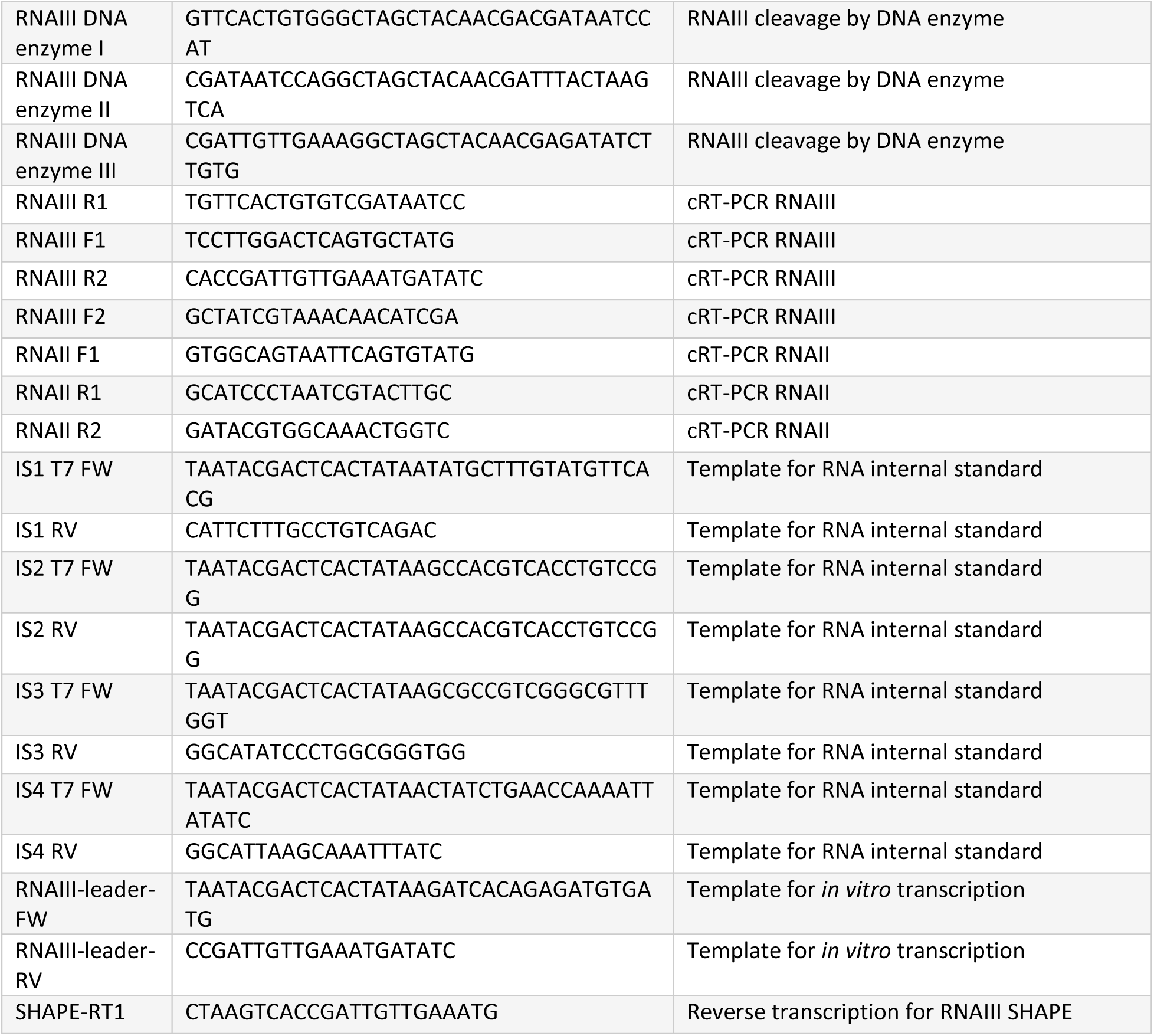
Oligonucleotides used in this study

**Table S4:**
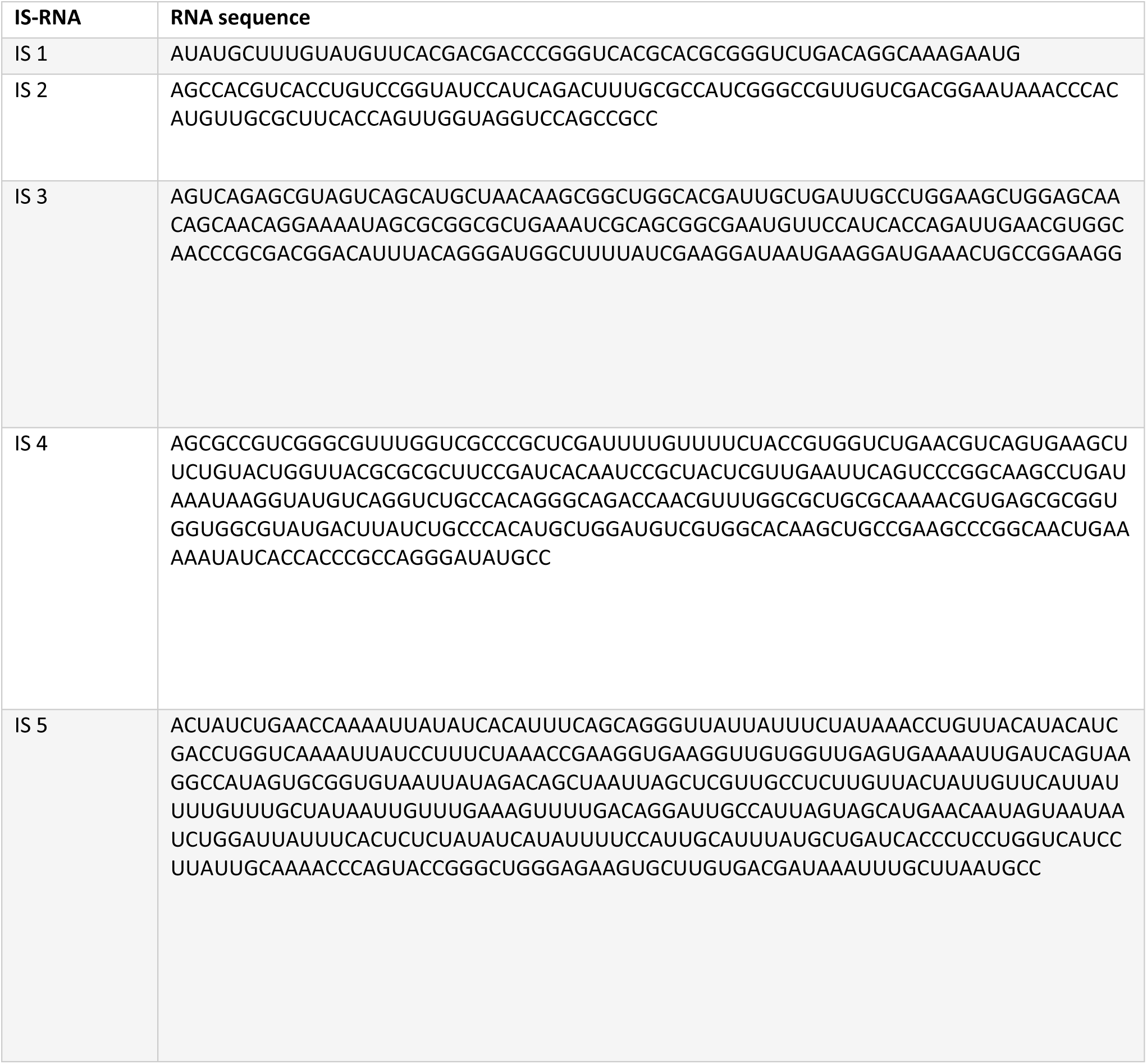
IS NAD-RNAs used for NAD captureSeq with their sequence.

**Table S5:** Enriched hits after NAD captureSeq in *S. aureus* pCG-P3 strain. Base mean: normalised mapped reads, Log2FC: Log2 Fold change (sample vs negative control). In bold are highlighted the common hits to *S. aureus* ATCC 25923 wild type strain (Table S2) and in regular font the non-common hits.

| Gene annotation number | Gene product | Base mean | Log2FC | P-value |
| --- | --- | --- | --- | --- |
| **SAOUHSC_00086** | **3-ketoacyl-acyl carrier protein reductase putative** | **60.65** | **2.49** | **0.000** |
| **SAOUHSC_00861** | **Lipoyl synthase** | **32.35** | **3.85** | **0.000** |
| **SAOUHSC_01685** | **Heat-inducible transcription repressor HrcA** | **127.01** | **2.84** | **0.000** |
| **SAOUHSC_02260** | **Delta-hemolysin precursor/RNAIII** | **5338.08** | **2.77** | **0.000** |
| SAOUHSC_00467 | Pur operon repressor | 24.92 | 2.12 | 0.000 |
| SAOUHSC_01191 | 50S ribosomal protein L28 | 18.24 | 2.28 | 0.000 |
| SAOUHSC_01432 | Peptide methionine sulfoxide reductase | 13.50 | 3.53 | 0.000 |
| SAOUHSC_01493 | 30S ribosomal protein S1 putative | 36.32 | 2.05 | 0.000 |
| SAOUHSC_01997 | Peroxide-responsive repressor PerR | 28.14 | 2.07 | 0.000 |
| SAOUHSC_02019 | Probable autolysin LytO | 203.90 | 2.52 | 0.000 |
| SAOUHSC_02409 | Arginase | 13.34 | 5.09 | 0.000 |

**Table S6:** Enriched hits after NAD captureSeq in *S. aureus* pCG-P3(−1G) strain. Base mean: normalised mapped reads, Log2FC: Log2 Fold change (sample vs. negative control). In bold are highlighted the common hits to *S. aureus* ATCC 25923 wild type strain (Table S2) and in regular font the non-common hits.

| Gene annotation number | Gene product | Base mean | Log2FC | P-value |
| --- | --- | --- | --- | --- |
| **SAOUHSC_00086** | **3-ketoacyl-acyl carrier protein reductase putative** | **300.15** | **3.06** | **0.000** |
| **SAOUHSC_00781** | **HPr kinase/phosphorylase** | **31.15** | **3.96** | **0.000** |
| **SAOUHSC_01176** | **Guanylate kinase** | **12.58** | **2.53** | **0.000** |
| **SAOUHSC_01617** | **Arginine repressor** | **22.34** | **2.28** | **0.000** |
| **SAOUHSC_01685** | **Heat-inducible transcription repressor HrcA** | **184.56** | **2.50** | **0.000** |
| **SAOUHSC_02260** | **Delta-hemolysin precursor** | **16912.85** | **3.57** | **0.000** |
| **SAOUHSC_02577** | **Putative 2-hydroxyacid dehydrogenase** | **66.55** | **2.07** | **0.000** |
| SAOUHSC_01191 | 50S ribosomal protein L28 | 67.43 | 2.27 | 0.000 |
| SAOUHSC_01653 | Superoxide dismutase 1 | 63.09 | 3.30 | 0.000 |
| SAOUHSC_01981 | Sensor histidine kinase putative | 16.93 | 5.07 | 0.000 |
| SAOUHSC_02019 | Probable autolysin LytO | 3110.21 | 3.23 | 0.000 |
| SAOUHSC_02409 | Arginase | 26.68 | 3.04 | 0.000 |

**Table S7:** Enriched hits after NAD captureSeq in *S. aureus* pCG-P2 strain. Base mean: normalised mapped reads, Log2FC: Log2 Fold change (sample vs negative control). In blue are highlighted the common hits to *S. aureus* ATCC 25923 wild type strain (Table S2) and in green the non-common hits. Note that *hld/*RNAIII gene is not enriched in this table because due to its +1G condition.

| Gene annotation number | Gene product | Base mean | Log2FC | P-value |
| --- | --- | --- | --- | --- |
| SAOUHSC_00086 | 3-ketoacyl-acyl carrier protein reductase putative | 47.53 | 2.68 | 0.000 |
| SAOUHSC_00861 | Lipoyl synthase | 16.34 | 3.30 | 0.000 |
| SAOUHSC_01685 | Heat-inducible transcription repressor HrcA | 71.22 | 2.81 | 0.000 |
| SAOUHSC_01191 | 50S ribosomal protein L28 | 17.10 | 2.11 | 0.000 |
| SAOUHSC_01997 | Peroxide-responsive repressor PerR | 18.60 | 2.19 | 0.000 |
| SAOUHSC_02019 | Probable autolysin LytO | 125.51 | 3.02 | 0.000 |

**Table S8:** pCG-P3 vs pCG-P3(−1G) transcriptomic analysis. Base mean: normalised mapped reads, Log2FC: Log2 Fold change (sample vs negative control). Negative Log2 Fold change values indicate downregulation, whereas positive values account for upregulated genes.

| Gene annotation number | Gene product | Base mean | Log2 Fold change | P-value |
| --- | --- | --- | --- | --- |
| SAOUHSC_00069 | Protein A | 94209.43 | -1.41 | 0.000 |
| SAOUHSC_01938 | Serine protease SplD | 502.55 | 1.06 | 0.000 |
| SAOUHSC_01939 | Serine protease SplC | 547.92 | 1.15 | 0.000 |
| SAOUHSC_01941 | Serine protease SplB | 584.74 | 1.19 | 0.000 |
| SAOUHSC_01942 | Serine protease SplA | 497.76 | 1.12 | 0.000 |
| SAOUHSC_02576 | Secretory antigen precursor SsaA putative | 560.25 | -1.28 | 0.000 |

**Table S9:**
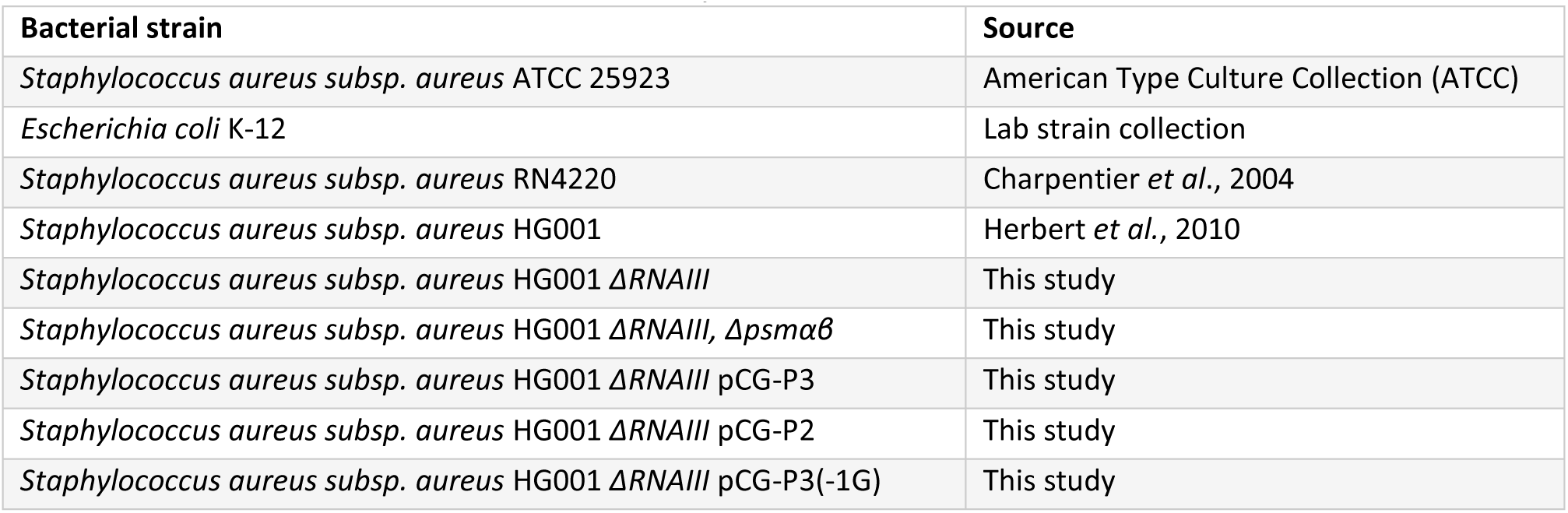
Bacterial strains used in this study

